# Scaled Recoupling of Chemical Shift Anisotropies at High Magnetic Fields under MAS with Interspersed *C*-elements

**DOI:** 10.1101/2020.07.02.184770

**Authors:** Keith J. Fritzsching, Eric G. Keeler, Chengming He, Ann E. McDermott

**Affiliations:** Department of Chemistry, Columbia University, New York, NY 10027, United States

## Abstract

The power of chemical shift anisotropy (CSA) measurements for probing structure and dynamics of molecules has been long recognized. NMR pulse sequences that allow measurement of CSA values in an indirect dimension of a protein correlation spectrum have been employed for aliphatic groups, but for practical reasons carbonyl functional groups have been little studied, despite the fact that carbonyls are expected to give particularly varied and informative CSA values. Specifically, the wide spectral widths of carbonyl tensors make their measurements difficult with typically attainable spectrometer settings. We present here an extended family of experiments that enable the recovery of static CSA lineshapes in a indirect dimension of magic angle spinning (MAS) solid-state NMR experiments, except for various real valued scaling factors. The experiment is suitable for uniformly labeled material, at moderate MAS rates (10-30 kHz), and at higher magnetic fields (*ν*_0*H*_ > 600 MHz). Specifically, the experiments are based on pulse sequence elements from a previous commonly used pulse sequence for CSA measurement, ROCSA, while modification of scaling factors is achieved by interspersing different blocks of *C*-elements of the same 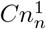 cycle. Using experimental conditions similar to the parent ROCSA sequence, a CSA scaling factor between 0 and 0.272 can be obtained, thus allowing a useful practical range of possibilities in experimental conditions for measurement of larger CSA values. Using these blocks it is also possible to make a constant-time CSA recoupling sequence. The utility and effectiveness of this approach, *f*ROCSA, is shown on model compounds 1-^13^C-Gly, U-^13^C,^15^N-l-His, and microcrystalline U-^13^C,^15^N-Ubiquitin.

## I. INTRODUCTION

The orientation dependence of the NMR chemical shift, specifically its range and distribution of values, referred to as the chemical shift anisotropy values, or CSA, reflect the molecular environment of the nuclear spin. CSA has been used as a powerful restraint on structure, including both conformation and non bonded interactions as well as dynamics. Interest in molecular origins of chemical shielding is essentially as old as NMR itself, and cross polarization techniques greatly enabled studies of the CSA of carbonyls and the influence of hydrogen bonding and other molecular influences.^1,2^ The relationship between CSA values for carbon atoms and structure in proteins is particularly noteworthy, since the CSA values provide probes of the local secondary structure,^3^ hydrogen bonding environment,^4,5^ and other longer range electronic effects.^6^ The trends and origins for aliphatic carbons have been well explored. Quantum chemical calculations, as well as bioinformatic tools, have been used to interpret CSA values for backbone C*α* carbons in proteins to unveil torsional states and secondary structure.^7,8^ Carbonyl and carboxylic acid CSA values are of particular interest; they can be used for understanding or determining protein structure and function, are sensitive to hydrogen bonding interactions, and also serve as sources of powerful orientational restraints.^5,6,9–11^ Additionally, the tensor values can provide the basis for probing protein dynamics. Fluctuations of the orientation of a CSA tensor as in chemical exchange processes also results in spin relaxation for solution^12,13^ as well as solid state NMR.^14^ Studying CSA re-orientation driven relaxation is promising for monitoring intermediate time-scale chemical exchange, for example by measuring relaxation during an applied spin locking field.^15^ These phenomena in aggregate provide strong motivation for measuring carbonyl CSA values.

Many solid-state NMR experiments on complex systems rely on magic-angle-spinning (MAS) to improve resolution,^16^ so it is of interest to measure anisotropic interactions during MAS. In recent work, a common class of experiments for accessing these interactions during MAS uses synchronous radio frequency (RF) pulses with the rotor period to partially recouple the anisotropic interactions.^17–26^ Other approaches to determine the tensor values during MAS include extracting information from the spinning sidebands,^27–33^ or using exotic hardware to vary the spinning axis over time.^34–38^ Ideally, the recoupling experiments are selective in the sense that the effective Hamiltonian has essentially only one specific interaction of interest, undistorted from its static form so that the lineshape is a scaled version of the familiar static lineshape, or alternatively produces a lineshape that can be uniquely simulated to extract the tensor parameters. In either case, it is desirable to produce a lineshape that is sensitive to all tensor parameters and not only the breadth. Typically pulse sequences for measuring anisotropic interactions are characterized by their scaling factors (the ratio of the apparent tensor width to its true width measured directly), and for many applications with relatively modest CSA spans it has been useful to identify sequences with the largest possible scaling factors so that the parameters of the interaction tensor are readily extracted. We examine the opposite situation in this work, namely an optimal sequence for measurement of tensors with unusually broad spans.

Recoupling the CSA during MAS was first shown in the seminal work of Alla *et al*.^17^ (2 *π*-pulse/*τ_r_*) and by Yarim-Agaev *et al*.^18^ (6 *π*-pulse/*τ_r_*). Subsequently, a variety of practical schemes to recouple the CSA have been reported. An experiment called Recoupling Of Chemical Shift Anisotropy, or ROCSA,^24^ based on symmetric pulse sequence elements^39^ has proven useful for measuring both ^13^C and ^15^N CSAs in uniformly isotopically enriched proteins.^7,40–42^ The two-dimensional and threedimensional ROCSA experiments are shown in Figure S1 (with *α*=0.0329 and *b*=0.467 and the A block indicated in Figure 1a). This pulse sequence has been extensively used to measure of the C*α* and other aliphatic CSA tensors, which have been shown to be useful for probing protein conformation. Meanwhile, it has been challenging to measure larger CSA tensor values using ROCSA due to common practical experimental limitations (the combination of available static applied field strengths, desirable or possible spinning frequencies consistent with stable experimental conditions, and limited RF field strengths). In particular, studies of the carbonyl group are hampered by this limitation as discussed further in the discussion section. Alternative pulse programs have also been developed to determine the CSA tensor parameters. One such approach uses *R*-symmetry elements to recouple the spatial CSA components and lineshapes derived with this experiment have been fit to measure the magnitude of the anisotropy (|*δ*_CSA_|) and the asymmetry (*η*_CSA_) of the CSA tensor.^26,43–47^ While this sequence has provided a solution for broad tensors,^48^ the *R*-symmetry sequences produce a symmetrized lineshape, from which the sign of the anisotropy cannot be determined and the asymmetry parameter is nearly indistinguishable when < 0.5.^49^

**FIG 1:**
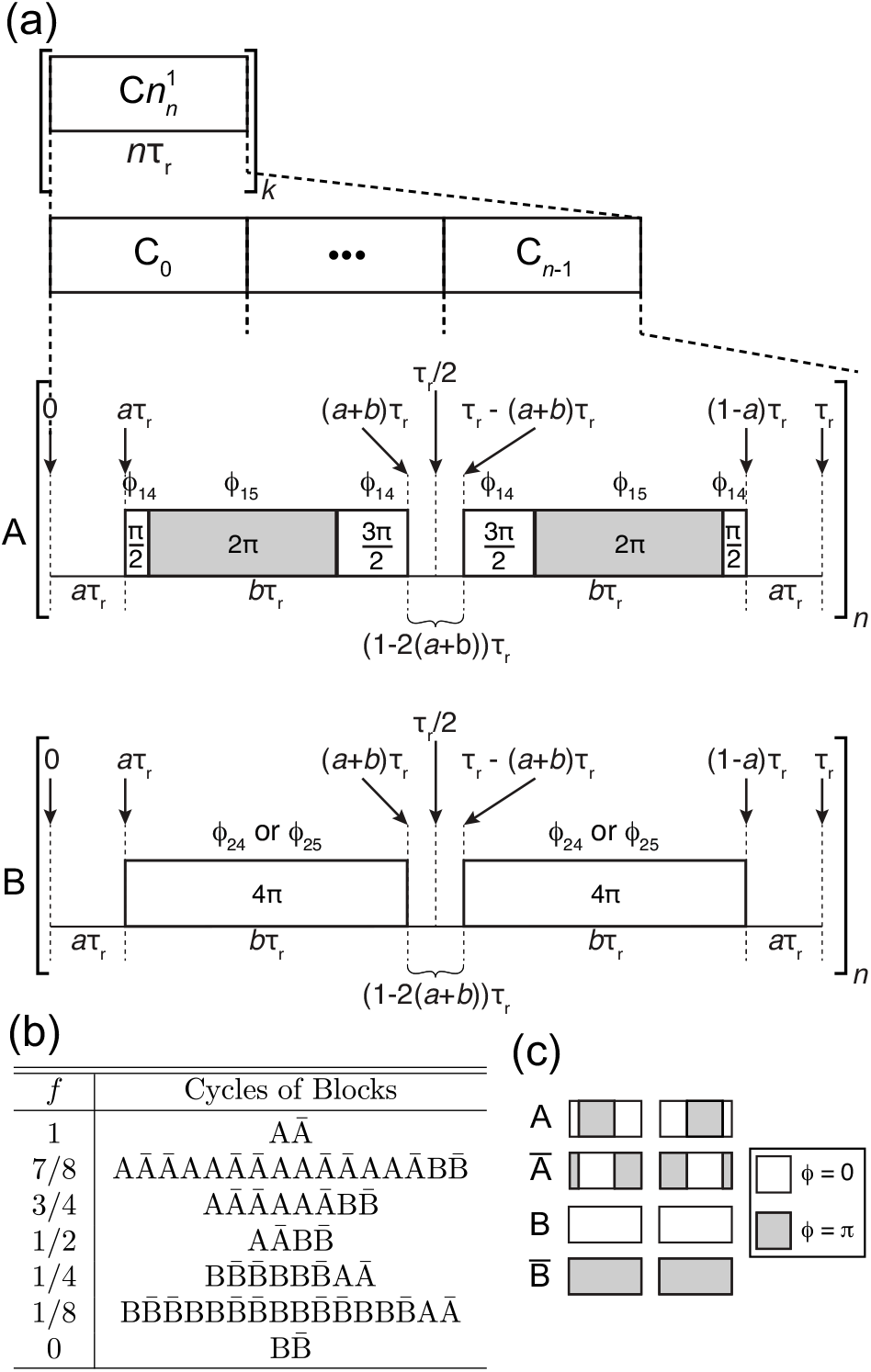
Pulse sequence diagram for *f*ROCSA. (a) The A and B blocks of *f*ROCSA depict the pulse timings and phases for two different *C*-symmetry blocks. The various *f*ROCSA pulse sequences are produced by interspersing A and B blocks to yield different real scaling factors, given in Table I. The RF field strength is *ν*_1_ = 2*ν_r_* = *b*. For the A block *b* = 0.467 and for the B block *b* = 0.467 or *b* = 0.5 (b) The versions of the *f*ROCSA experiments that are demonstrated in this study. The cycles are defined as a combination of blocks (A and B) and phase inverted blocks (Ā and 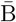). The length of a cycle is determined by 2x the denominator of the fraction, *f*. (c) A single element of the A and B blocks are shown with their respective phase differences. For 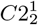 the blocks are phase cycled according to: *ϕ*_14_ 0220, *ϕ*_15_ = 2002, *ϕ*_24_ = 2002, *ϕ*_25_ = 0220. In experiments where both A and B blocks are used, the notation *a*_1_, *b*_1_ and *a*_2_, *b*_2_ indicates the *a*, *b* values for the A and B blocks, respectively, *e.g*., *f*ROCSA(*a*_2_, *b*_2_).

Motivated by improving precision of measurements of larger CSA values, we present a family of experiments, called *f*ROCSA for fractionally scaled ROCSA, that, like the parent ROCSA experiment,^24^ preserve the CSA powder lineshape. The f ROCSA experiments in addition allow investigator selection of the scaling factor in a way that allows for CSA with larger spans to be studied at moderate spinning frequencies (*ν_r_* = 10-30 kHz), realistic RF field strengths (*ν*_1_ = 2*ν_r_*/*b* ≈ 4*ν_r_*), and high external static magnetic fields (*ν*_0*H*_ > 600 MHz).

## II. THEORY

Using approaches similar to those used in the development of ROCSA^24^ we identified pulse sequences that, like ROCSA, recover the static CSA lineshape and limit the effect of homonuclear dipolar coupling, but in addition have variable scaling factors to accommodate a range of CSA tensor widths. Specifically, we identified additional *C*-elements that for 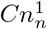 symmetric sequences result in different real valued scaling factors. By judiciously averaging these *C*-elements over the *k* repeated blocks and over the *s* different scans the effective average Hamiltonian can be tuned, thus creating a larger family of sequences and expanding the possibilities for the original ROCSA sequence.

### A. Average Hamiltonian

Throughout we use notation and formalism similar to previous work,^39,50,51^ where the high-field truncated Hamilton is represented:

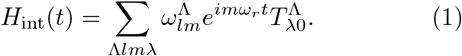

Here Λ represents a specific interaction (*e.g*., CSA, dipolar coupling), *l* represents the spatial rank, *m* is the spatial quantum number, λ is the spin rank, *μ* is the spin quantum number, *ω_r_* = 2*π*/*τ_r_* is the rotation frequency around the magic angle (where 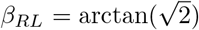), and an arbitrary initial rotor phase angle *α_RL_* has been assumed. The characteristic frequency 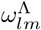 for the spatial tensor components 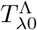 is expressed in terms of complex amplitudes 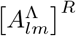 in the rotor frame *R* (with respect to the lab frame *L*):

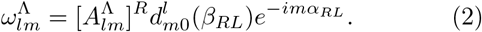

Interactions with spatial rank *l* = 2, such as the CSA or dipolar coupling, have zero amplitude for the component with spatial quantum numbers of zero *m* = 0, since the Wigner element 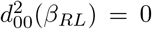 during MAS. The rotor frame is related to the principle frame, *P*, for the interactions Λ through the molecular frame, *M*, by:

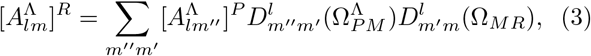

where, 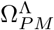 and 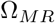 are sets of Euler Angles describing the transformations between the different frames.

The ROCSA sequence, and this modification *f*ROCSA, is constructed within the symmetry class 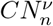. *C* refers to any RF element that performs an integer number of 2*π* rotations (including windowed examples). Such a *C*-element, as demonstrated for 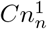 in Figure 1a, is repeated *N* times in *n* rotor periods with *ν*, the spin winding number, indicating the RF phase shift of the *q*th *C*-element by 2*πqν*/*N*, where *q* is the integer from 0…*N*-1.^52^ If the system is stroboscopically sampled at integer numbers of the rotation period *τ_r_*, the first-order average Hamiltonian of pulse sequences in this class in the RF toggling frame is:

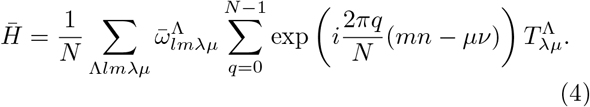

Here 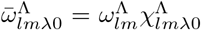, meaning that the effective amplitude for the tensor components during the sequence are related to the static values defined above by complex scaling factors with magnitudes < 1, designated 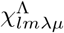.

For pulse sequences with 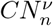 symmetry, the first-order average Hamiltonian has the selection rule,^50^

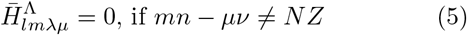

where *Z* is an integer including 0. If the pulse sequence has 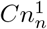 symmetry, the relation is than simplified to:

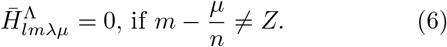

If *n* = 1, all terms are symmetry allowed. If *n* = 2, the CSA terms with quantum numbers {*l*; *m*; *λ*; *μ*} = {2; ±1, ±2;1;0} and the dipolar terms {2; ±1, ±2; 2; 0, ±2} are symmetry allowed. If *n* ≥ 3 only terms with *μ* = 0 are symmetry allowed. Regardless of *n*, pulse sequences with 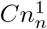 symmetry have an effective Hamiltonian:

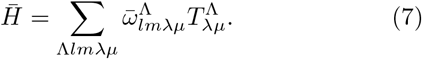

If the *μ* ≠ 0 terms are zero, the effective Hamiltonian has the same form as the static Hamiltonian (*i.e*., setting *ω_r_* = 0 in Eq. 1).

### B. Scaling Factors

To explore how the scaling factors depend on the time-dependent RF pulses it is cumbersome, but useful, to write the RF rotations of the *C*-element with a set of time-dependent Euler angles, Ω_rf_(*t*) = {*α*_rf_(*t*), *β*_rf_(*t*), *γ*_rf_(*t*)}. In the simple case of amplitude modulated pulses:

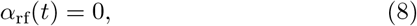

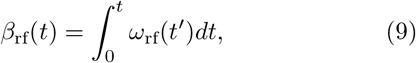

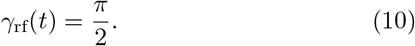

*ω*_rf_(*t*) is the effective RF field strength. A negative value for the field strength was used to represent 180° phase shifted pulses. *β*_rf_ is calculated by integrating piece-wise over the segments of the pulse sequence element. For 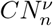 the scaling factors can be expressed:

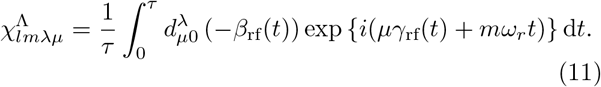

The present application is to identify pulse sequences that reproduce the static lineshape of the CSA while removing homonuclear dipolar terms. (It is assumed that heteronuclear dipolar terms would be removed by decoupling). To remove homonuclear dipolar couplings and scale the CSA flexibly, a pair or group of *C*-elements were sought such that for each element 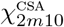 are identical real numbers for *m* = ±1, ±2. It is also required that the 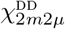 scaling factors are approximately zero but not necessarily identical for *m* = ±1, ±2. Real valued scaling factors are obtained for *C*-elements with symmetry imposed:

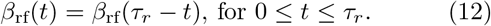

In this paper we refer to elements that satisfy this property as symmetric *C*-elements. To minimize 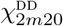 and vary 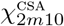 we searched the parameter space of *β*_rf_(*t*).

*C*-elements can be interspersed through out the sequence and over different scans to change the effective scaling factor. The effective scaling factor is then simply the weighted average of the different scaling factors:

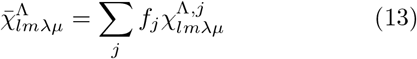

where the sum is over the different types of elements labeled by *j*, and *f_j_* fraction of such blocks (i,e, 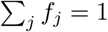)

To fully describe the pulse windows in the *C*-elements we define *a* and *b*, where *aτ_r_* is the length before pulsing starts and *bτ_r_* is the duration of the pulse, analogously to Chan and Tycko^24^.

#### 1. A-element

The original ROCSA sequence *C*-element is built using two POST composite pulses. POST pulse sequence elements are designed to be tolerant to RF inhomogeneity,^53^ and generate an effective 0*π* rotation. For the A-element as shown in Fig. 1a, 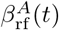 =

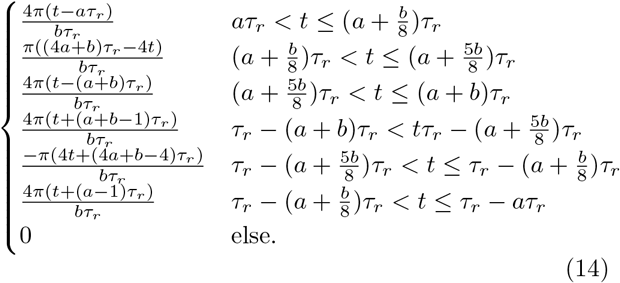

This function is integrated using Eq. 11 under the assumptions: 0 < *a* < 1/2, 0 < *b* < 1/2, and *a* + *b* < 1/2 to yield the scaling factors as a function of *a* and *b*:

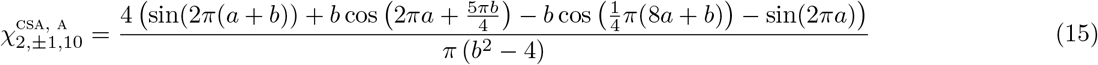

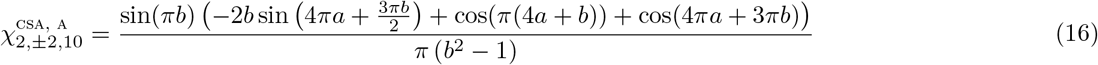

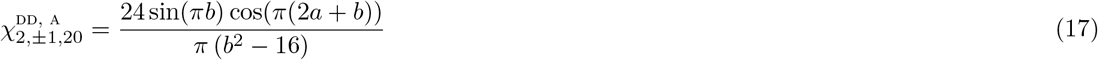

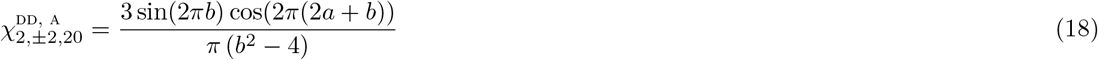

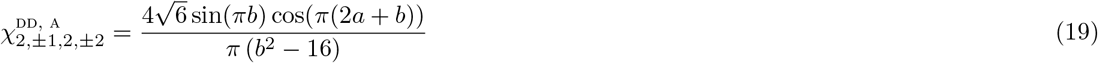

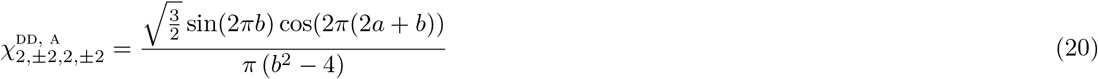

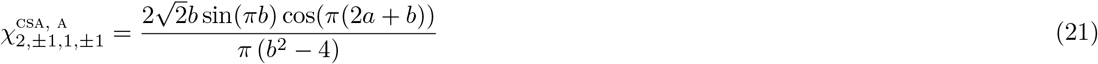

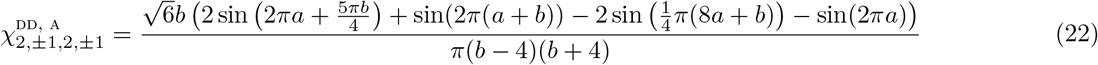

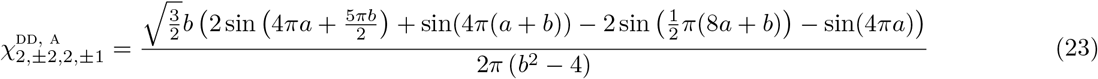

If *n* ≥ 3 only Eq. 15–18 need to be considered. If *n* = 2, the *μ* = ±2 terms additionally contribute (Eq. 19–20). As previously reported for ROCSA (*a*=0.0329, *b*=0.467) the *m* = ±2, *μ* = ±2 terms are small (≤ 0.0208). While the *n* = 1 condition to our knowledge was not previously discussed, it would be a potentially useful experiment since it maximizes spectral width. Unfortunately though for n=1 all of the terms contribute. The 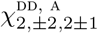 terms at the ROCSA condition are most problematic with a scaling factor of 0.0923; this is approximately double the scaling factor of the homonuclear scaling factors in the *n* ≥ 2 versions of ROCSA.

**FIG. 2.**
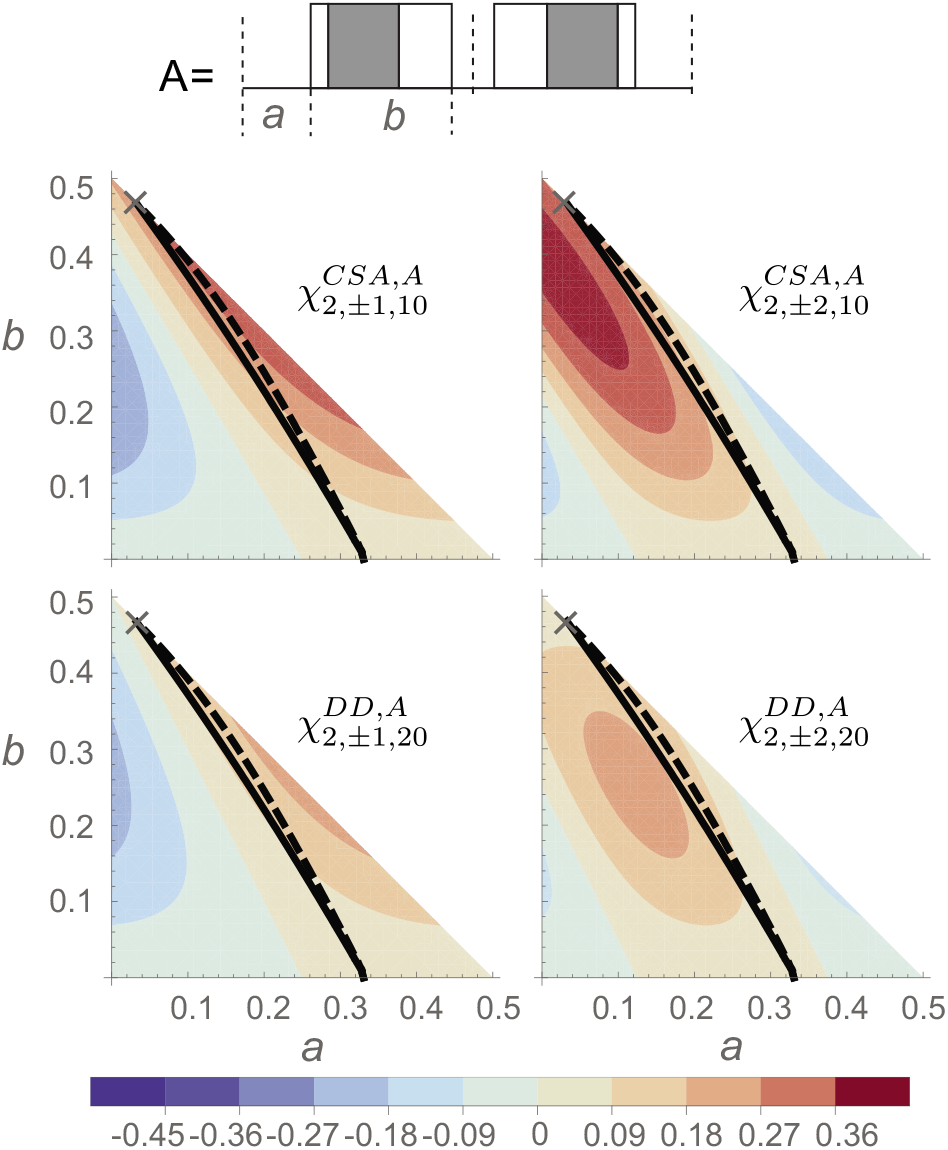
Contour plots of the scaling factors from Eq. 15–18 (A block). The solid line shows where 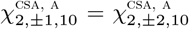. The dashed line shows were 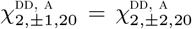. The additional scaling factors (Eqs. 19–23) are not shown. Gray Xs mark the (*a, b*) combinations that are used in this work.

It is noted that if the the POST pulses are reversed in time, *i.e*., so that the first pulse in the cycle is (3*π*/2)_*ϕ*_(2*π*)_*ϕ+π*_(*π*/2)_*ϕ*_ (instead of (*π*/2)_*ϕ*_(2*π*)_*ϕ+π*_(3*π*/2)_*ϕ*_) and Eq. 12 is still made valid, Eq. 14 is changed and the scaling factors are not the same. This pulse sequence would have a lower CSA scaling factors but the same dipolar scaling factors. It would also require higher RF field strengths to recover the static lineshape.

#### 2. B-element

Next we consider the simpler B-element, consisting of two 4*π* pulses as shown in Fig. 1a, 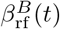 =

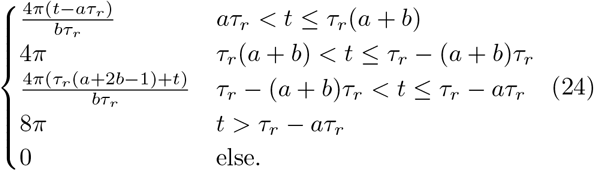

This function is likewise integrated using Eq. 11 under the same assumptions as before to generate another set of scaling factors:

**FIG. 3.**
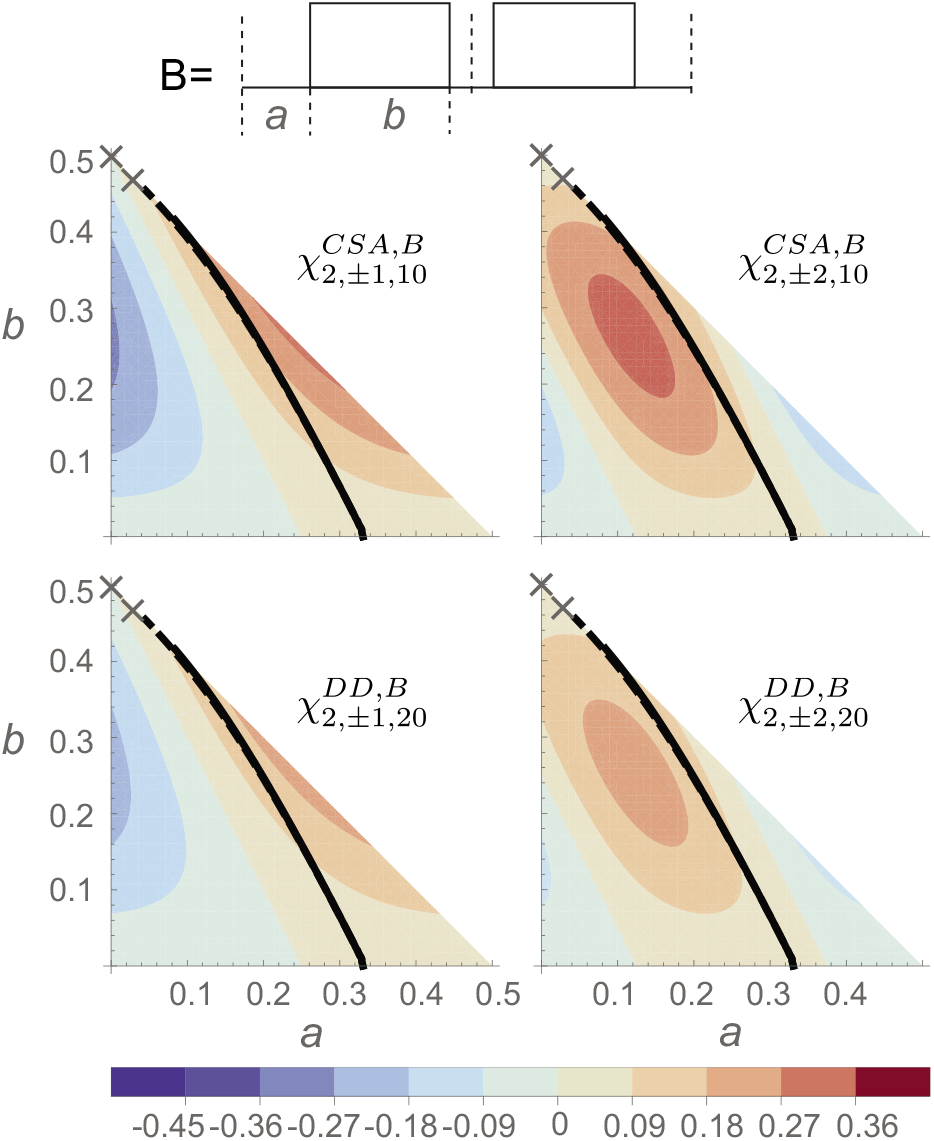
Contour plots of the scaling factors from Eq. 25–28 (B block). The solid line shows where 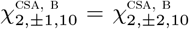. The dashed line shows were 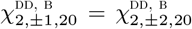. The additional scaling factors (Eqs. 29–30) are not shown. Gray Xs mark the (*a, b*) combinations that are used in this work.

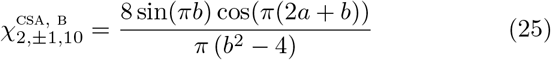

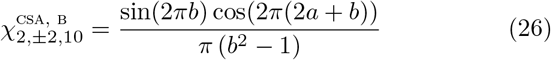

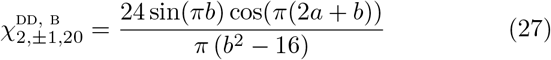

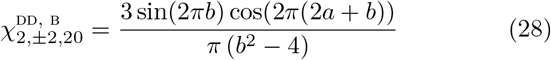

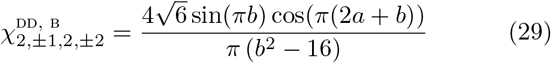

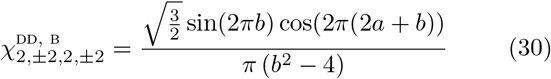

Similarly, for *n* ≥ 3 only the *μ* = 0 terms are symmetry allowed. The remainder of the terms are needed for *n* = 2. We did not consider the element for *n* = 1. In the limit: 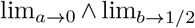 the symmetry of the B-elements can be recast as 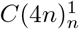. For these parameters, and for *n* ≥ 3 only the isotropic chemical shift and j-coupling terms are allowed. For *n* = 2, the dipolar terms 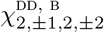 are also symmetry allowed, but have a scaling factor of zero.

#### 3. B’-element

Lastly, we introduce one additional block called B’, again consisting of two 4*π* pulses but with a 180 degree phase shift after each 2*π* rotation (shown in Figure S2). Using an identical approach to above, it can be shown that for *n* ≥ 2 the block has the the same scaling factors as the B-block (Eq. 25–30). As shown in the Supporting Information, B’ is more robust to frequency offset than B.

Therefore, we have identified different blocks, A and B (and B’), that enable us to tune the scaling factors.

### C. Constant-time

Constant-time pulse sequences can be used to decrease unwanted line broadening caused by other evolving couplings and relaxation processes in recoupling experiments. As was illustrated for example in PITHIRDS,^54^ zero scaling elements can be useful for generating constant-time recoupling sequences. A constant time sequence is split into effective blocks of time *τ_e_* and ineffective recoupling blocks of time *τ*_0_. The total recoupling time *t* = *τ_e_* + *τ*_0_ is fixed while *τ_e_* is incremented and *τ*_0_ is decremented correspondingly.

More generally, fractional symmetry element based constant-time experiments can be understood as operating by incrementing the scaling factors as the indirect dimension progresses (instead of incrementing time). This approach has been used before in another CSA recoupling experiment and was realized by changing the positions of the pulses in each block.^25^ Here we can simply increment 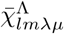 by changing the fraction *f_j_* of effective blocks in each indirect dimension increment. It is then feasible to increment in a way that gives CT-ROCSA or more generally CT-*f*ROCSA. The constant-time experiments have the same CSA scaling factors as the other experiments, and require no extra phase cycling. The only penalty is that all increments are subject to the same relaxation, and this results in a reduction in signal-to-noise.

## III. RESULTS

The *f*ROCSA based pulse sequences for carrying out two-dimensional CP-f ROCSA and three-dimensional DCP-f ROCSA experiments used in this study and shown in Figure S1 along with the required phase cycles. Two distinct types of *C*-symmetry blocks are interspersed in the pulse sequences, referred to as A and B blocks, and shown in detail in Figure 1. We refer to this experiment as *f*ROCSA, for fractionally scaled ROCSA, where *f* indicates the fraction of A blocks in the sequence. Specifically, the *f*ROCSA pulse sequence is composed of elements, blocks, and cycles. We put *n* elements in each block. The number of blocks in a cycle is equal to 2x the denominator of *f*, and is composed of blocks (A and B) and phase inverted blocks (Ā and 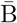). The phase inverted blocks are depicted with shading of the RF pulses in Figure 1c. We iterate the blocks and phase inverted blocks through the cycle. This cycle is continued through both the *k* points of the indirect dimension (*t*_1_ increments) and *s* scans. To ensure that the cycle is completed in each point of the indirect dimension, the number of scans, *s*, must be an integer multiple of 2x the denominator of *f*. The A-element is composed of two POST pulses, which is the same as in ROCSA, with timings of *a*_1_=0.0329 and *b*_1_=0.467. The B Block has *n* B-elements composed of two 4*π* pulses with timings defined by values *a*_2_ and *b*_2_. More generally we use the nomenclature *f*ROCSA(*a*_2_, *b*_2_), to account for the variable timing in the B-elements.

Experimental results for various *f*ROCSA(*a*_2_, *b*_2_) experiments are shown for the carboxyl and carbonyl carbons of 1-^13^*C*-glycine, U-^13^C,^15^N-l-histidine, and U-^13^C,^15^N-Ubiquitin. The CSA tensor parameters for the carboxyl and carbonyl carbons in these samples are extracted from each of the various *f*ROCSA(*a*_2_, *b*_2_) sequences implemented.

### A. Glycine

We characterized the CSA values for 1-^13^C-glycine in the *α* polymorph by non-spinning, ROCSA, and *f*ROCSA experiments. We compared CSA tensor parameters determined in a non-spinning experiment (*δ*_CSA_ = −71.1 ± 0.7 ppm and *η*_CSA_ = 0.91 ± 0.01) to those determined by various *f*ROCSA(*a*_2_, *b*_2_) experiments (Figure S3 and Table SI). The experimental scaling factors for the *f*ROCSA(*a*_2_, *b*_2_) experiments were also determined using the CSA tensor parameters from non-spinning data. The CSA tensor parameters obtained in this study from all techniques agree with previously reported values,^55,56^ with deviations that are well within the experimental error, offering strong support for the accuracy of the *f*ROCSA(*a*_2_, *b*_2_) experiments for determination of CSA tensor parameters.

### B. Histidine

The carboxyl carbon of U-^13^C,^15^N-l-histidine in the *τ*-tautomer form^57^ was used as a model to demonstrate the *f*ROCSA experiments and the ability to determine accurate CSA tensor parameters from the scaled CSA lineshape, as shown in Figures 4–5, S4-5, and Table I. The experimentally determined scaling factors and subsequent errors were determined by fitting the CSA lineshape using the best fit *δ*_CSA_ and *η*_CSA_ from the ROCSA experiment. Additionally, we verified the scaling factors and their experimental errors by performing the same fitting routine but replacing the *δ*_CSA_ and *η*_CSA_ with the best fit parameters for each *f*ROCSA experiment. There was little difference in the scaling factors and errors determined by these two methods and the former are shown in Figure 5. Moreover, excellent agreement between the theoretical scaling factors and those determined experimentally was observed (Table I). As shown in the theory section to first order and when *n* ≥ 2 the B block with *b* = 0.5 strongly attenuates rank two interactions; this is shown experimentally when only B blocks with *b* = 0.5 are used (Figures 4b,c and S6). This is a phase alternating spin lock with *ν*_1_ = 4*ν_r_*. Additionally, the CSA tensor parameters for the N^*δ*^ and N^*ϵ*^ nitrogens of the sample were also determined and are shown in Figure S7.

**FIG. 4.**
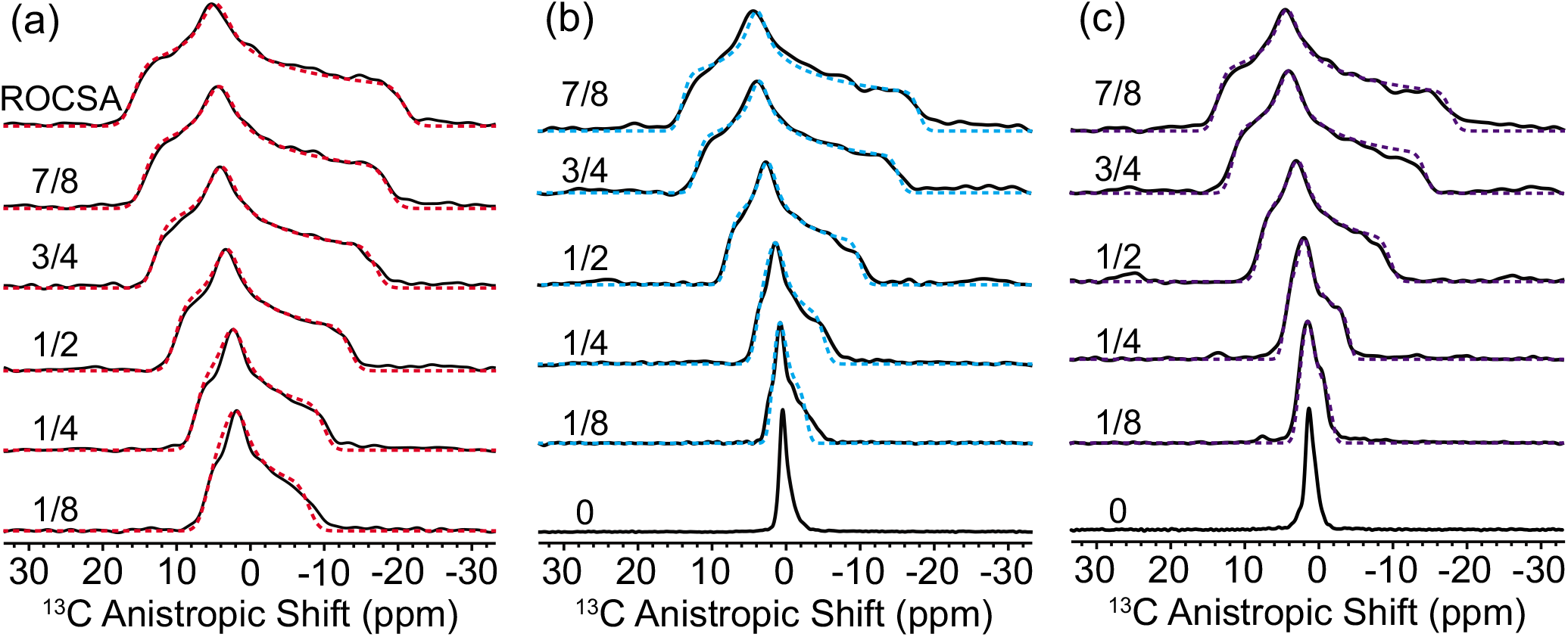
(a) 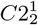 *f*ROCSA(0.0329,0.467) – B, (b) 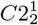 *f*ROCSA(0,0.5) – B, and (c) 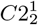 *f*ROCSA(0,0.5) – B’ experimental (solid, black) and simulated (dashed, red/cyan/purple) lineshapes of the carboxyl carbon of l-histidine for the various sequences (7/8 ≥ *f* ≥ 1/8) described in Figure 1. The *f*ROCSA(*a*_2_, *b*_2_) spectra are presented with the scaled anisotropic shift axis that is detected and not corrected for the scaling factor 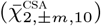. Simulated CSA tensor parameters for each experiment are given in Table I.

**FIG. 5.**
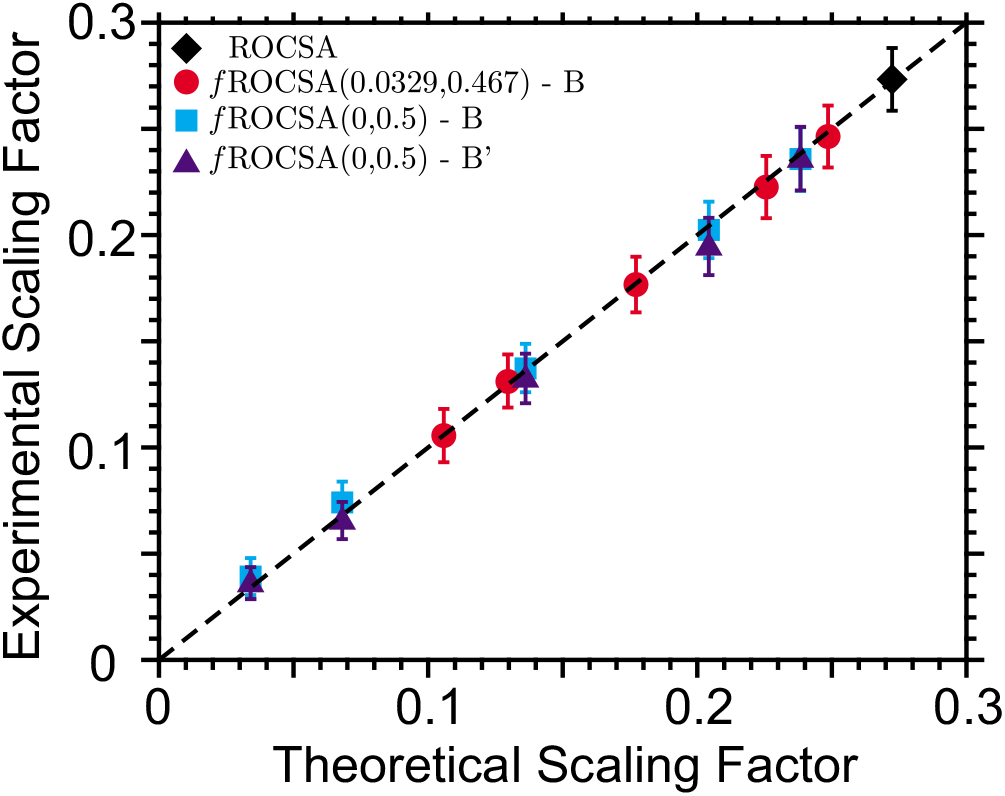
Plot of the theoretical and experimental CSA scaling factors 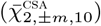 for ROCSA and the various *f*ROCSA(*a*_2_, *b*_2_) sequences. ROCSA is shown as a black diamond. The red circles represent the ROCSA and *f*ROCSA(0.0329,0.467) experiments, the cyan squares represent the *f*ROCSA(0,0.5) experiments that are run with the B block, and the purple triangles represent the *f*ROCSA(0,0.5) experiments run with B’ block.

**TABLE I.** Experimentally determined CSA tensor values and scaling factors from the carboxyl carbon of l-histidine

| Experiment | $\delta_{\text{CSA}}$ (ppm) | $\eta_{\text{CSA}}$ | Theoretical $\bar{\chi}_{2\pm m10}^{\text{CSA}}$<br>Scaling Factor | Experimental $\bar{\chi}_{2\pm m10}^{\text{CSA}}$<br>Scaling Factor<sup>a</sup> |
| --- | --- | --- | --- | --- |
| ROCSA | $-76 \pm 4$ | $0.54 \pm 0.08$ | 0.272 | $0.27 \pm 0.01$ |
| $f\text{ROCSA}(0.0329, 0.467) - \text{B}$ | | | | |
| 7/8 | $-75 \pm 5$ | $0.55 \pm 0.08$ | 0.249 | $0.25 \pm 0.01$ |
| 3/4 | $-75 \pm 5$ | $0.53 \pm 0.09$ | 0.226 | $0.22 \pm 0.01$ |
| 1/2 | $-76 \pm 6$ | $0.53 \pm 0.11$ | 0.177 | $0.18 \pm 0.01$ |
| 1/4 | $-76 \pm 8$ | $0.52 \pm 0.16$ | 0.130 | $0.13 \pm 0.01$ |
| 1/8 | $-75 \pm 9$ | $0.49 \pm 0.22$ | 0.106 | $0.11 \pm 0.01$ |
| $f\text{ROCSA}(0, 0.5) - \text{B}$ | | | | |
| 7/8 | $-75 \pm 8$ | $0.57 \pm 0.12$ | 0.238 | $0.24 \pm 0.01$ |
| 3/4 | $-75 \pm 8$ | $0.55 \pm 0.13$ | 0.204 | $0.20 \pm 0.01$ |
| 1/2 | $-74 \pm 10$ | $0.53 \pm 0.18$ | 0.136 | $0.14 \pm 0.01$ |
| 1/4 | $-77 \pm 15$ | $0.47 \pm 0.28$ | 0.068 | $0.07 \pm 0.01$ |
| 1/8 | $-81 \pm 16$ | $0.42 \pm 0.29$ | 0.034 | $0.04 \pm 0.01$ |
| $f\text{ROCSA}(0, 0.5) - \text{B}'$ | | | | |
| 7/8 | $-75 \pm 5$ | $0.53 \pm 0.08$ | 0.238 | $0.24 \pm 0.01$ |
| 3/4 | $-73 \pm 5$ | $0.52 \pm 0.09$ | 0.204 | $0.19 \pm 0.01$ |
| 1/2 | $-74 \pm 7$ | $0.51 \pm 0.13$ | 0.136 | $0.13 \pm 0.01$ |
| 1/4 | $-73 \pm 10$ | $0.55 \pm 0.20$ | 0.068 | $0.066 \pm 0.009$ |
| 1/8 | $-77 \pm 14$ | $0.54 \pm 0.30$ | 0.034 | $0.036 \pm 0.008$ |

<sup>a</sup> Experimental scaling factors were determined by the best fit of scaling factor parameter using the best fit parameters from the ROCSA experiment.

### C. Ubiquitin

To demonstrate the advantages of the *f*ROCSA(*a*_2_, *b*_2_) experiment for characterizing more complex systems we developed a three-dimensional NCO-1/2-ROCSA(0.0329,0.467) experiment and applied it to microcrystalline Ubiquitin. We measured site-specific CSA tensor parameters for the backbone carbonyl carbons in the uniformly ^13^C,^15^N enriched protein sample using the *f*ROCSA(*a*_2_, *b*_2_) experiment (Figures 6 and S8-10). While CSA tensor values of the carbonyl carbons of Ubiquitin have been determined using solution state NMR previously,^13,58–61^ to our knowledge they have not been previously reported based on solid-state NMR experiments.

**FIG. 6.**
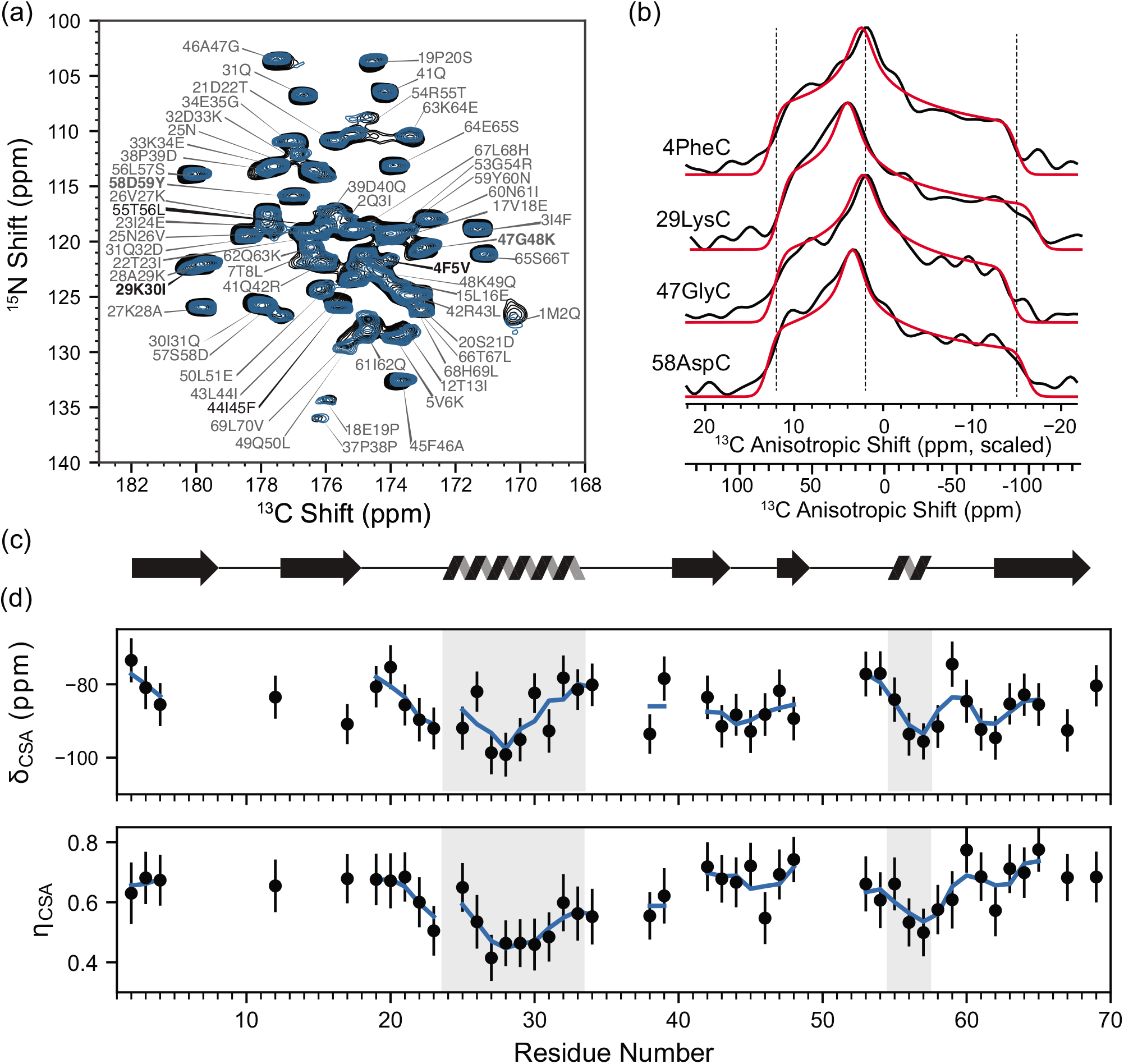
(a) Isotropic projection of 3D 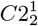 NCO-1/2-ROCSA(0.0329,0.467) (black) and 2D NCO (teal) spectra of microcrystalline U-^13^C, ^15^N-Ubiquitin. (b) Anisotropic projections of several peaks in (a) with best fit curves (red). The 1/2-ROCSA(0.0329,0467) experiment has a scaling factor 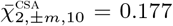 which is used to extract the CSA tensor parameters, both the scaled and unscaled anisotropic shift axes are shown. (c) Cartoon representation of the secondary structure of Ubiquitin. (d) The variation of *δ*_CSA_ (top) and *η*_CSA_ (bottom) for each resolved and fit CO atom by residue. The blue curve is the value averaged with its residue neighbors. Gray boxes indicate the location of *α* helices in the Ubiquitin structure.

## IV. DISCUSSION

The spectral widths of the indirect dimensions of the ROCSA and *f*ROCSA experiments (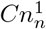 elements) are *ν_r_/n*, where *ν_r_* = *ω_r_*/2*π* is the MAS frequency in Hz. This limitation of the spectral width can be thought of as imposing a maximal external static magnetic field for the study of carbonyl chemical shift tensors by the ROCSA experiment of ~22 T (~940 MHz for ^1^H) if moderate spinning frequencies (10-20 kHz) and consequently moderate RF fields are used (*ν*_1_ = 2*ν_r_/b* or approximately 4 times the spinning frequency as *b* = 0.467 for the ROCSA experiment), as discussed previously.^24^ When considering ROCSA measurements for proteins or other biological systems additional practical restrictions may apply. Typically, spinning frequencies are chosen to avoid unwanted interference from rotational resonance (*R*^2^) conditions (*e.g*., ~85 ppm or >160 ppm). When selecting the moderate spinning frequency of ~85 ppm, the effective spectral width is given by 85 ppm/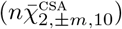 where the 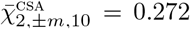 and *n*=2 for the typical ROCSA sequence (for carbonyls). This imposes a limitation on the measurement of CSA lineshapes such that the full breadth of the tensor Ω_CSA_ should be well below the effective sweep width of 156 ppm, and precludes the study of relatively large CSA tensors. If the effective spectral width would be near or less than the breadth of the CSA lineshape the ability to extract accurate tensor measurements is difficult and distorted lineshapes result (Figure S11). This consideration has inhibited the study of large CSA environments such as the protein carbonyl using the ROCSA experiment. If, on the other hand, the spinning frequency is >160 ppm, the spectral width of the experiment no longer limits the ability to study large CSA tensors with ROCSA. However, another consideration then arises with respect to common RF power limitations. The RF field strength required for the ROCSA experiment is *ν*_1_ = 2*ν_r_/b*, where *b* = 0.467, so for >160 ppm spinning frequencies requires *ν*_1_ > 685 ppm (>103 kHz if *B*_0_ > 600 MHz). Clearly as the applied field strength becomes higher this requirement becomes prohibitive. Another way to approach this issue is to examine the typical RF field strengths that are allowable for typical hardware. In this case, the maximally allowed spinning frequency can be defined by *ν_r,max_* = *b*_*ν*_1_,*max*_/2, where *ν*_1,*max*_ is the maximally allowed RF field strength for a particular hardware. When considering the typical experimental conditions (3.2mm probe with < 70 kHz or 1.9mm probe with < 90 kHz maximally allowed RF field strengths) there are few allowable ROCSA experimental conditions for the measurement of large CSA tensors (Ω_CSA_ > 156 ppm) that also avoid R^2^ conditions.

An alternative opportunity would be to use a lower symmetry element, 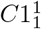, for the ROCSA experiment (Figure S12) which allows for the measurement of larger CSA tensors (*e.g*., 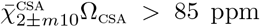) because it effectively doubles the sweep width of the indirect dimension. Unfortunately, as shown in the Theory section, for *n* = 1, 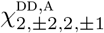 is not negligible, and may lead to distortion the static lineshape. As the most problematic scaling factor that is introduced when using *n* = 1 is due to homonuclear dipolar couplings, if these couplings are small, or non-existent, (*e.g*., sparsely ^13^C labeled samples) then the use of the 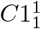 ROCSA or *f*ROCSA experiments with little distortion of the static lineshape is possible. Importantly, the experimental time for this experiments is doubled for acquiring the same indirect resolution. Lastly, the signal is spread out over a broader range in ROCSA than *f*ROCSA so the signal-to-noise is worse (see Figure S13), although there is also a trade off in preciseness. Overall 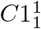 may be less practical than the fractional ROCSA options described here. We discuss how to determine the allowable experimental conditions in more detail using the example of our U-^13^C, ^15^N-Ubiquitin measurements in Figure S14. Another consideration is that when using the *f*ROCSA(*a*_2_, *b*_2_) experiments with small scaling factors the *n* > 2 *C*-symmetry elements can be used to measure larger CSA tensor environments (Figure S15, 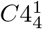 *f*ROCSA(0,0.5)). Constant time versions are also possible (Theory Section C), here we do not see much improvement in lineshape (Figure S16).

As discussed in Chan and Tycko^24^ the RF inhomogeneity of the ROCSA sequence must be considered for applications on complex systems. The POST pulses that are used in the ROCSA experiment and the A block of each *f*ROCSA variation presented here are designed to be tolerant to RF inhomogeneity.^53^ However, the 4*π* pulses that are used in the B block of *f*ROCSA are not as tolerant to RF inhomogeneity and distortions to the static CSA lineshape are larger. To demonstrate the effects of RF inhomogeneity on the experiments the RF field strengths were intentionally miscalibrated (Δ*ν*_1_±5,10 kHz, Figure S17 and Tables SIII-SV). These results show that the RF field strength for the *f*ROCSA periods must be carefully calibrated for each sample to ensure that the proper CSA lineshape is produced. Additionally, replacing the simple 4*π* pulse of the B block with a 2*π*(0)2*π*(*π*) pulse, referred to as B’, reduces the effects of *ν*_1_ miscalibration (Figures 4 and S2, and Tables I and SV). The scaling factors for the B’ block are unchanged from the B block. When RF inhomogeneity and miscalibration are thought to be problematic we recommend that the B’ block is used for *f*ROCSA experiments to reduce these effects.

## V. CONCLUSIONS

A modified version of ROCSA, called *f*ROCSA, is presented and shown to have the ability to probe environments with large chemical shift anisotropies in complex systems where site-specific resolution is required. The *f*ROCSA experiment operates by averaging different blocks of the same *C*-element over *k* repeated blocks of an indirect dimension and *s* different scans. In this way the effective average Hamiltonian can be adjusted allowing for user selection of the real valued scaling factors to produce an undistorted scaled CSA lineshape. We show that this sequence is applicable to moderate spinning frequencies (*ν_r_* = 10-30 kHz) and high external static magnetic fields (*ν*_0*H*_ > 600 MHz) where many characteristic experiments on complex systems, in particular studies of peptides and proteins, are performed. We determined the CSA tensor parameters of the carboxyl carbon for crystalline 1-^13^*C*-glycine, and U-^13^C, ^15^N-l-histidine using multiple different variations of *f*ROCSA(*a*_2_, *b*_2_).

These initial experiments on model systems verify the ability to select the real valued scaling factors (between 0 and 0.272) of the *f*ROCSA family of pulse sequences to generate a scaled static CSA lineshape. Comparisons with previous literature values and across the *f*ROCSA landscape (7/8 ≥ *f* ≥ 1/8) demonstrate the ability to faithfully extract the CSA tensor values from the scaled spectrum. Furthermore, we demonstrate the use of *f*ROCSA(a_2_,b_2_) on a complex system, microcrystalline U-^13^C, ^15^N-Ubiquitin, to extract the CSA tensor parameters of the backbone carbonyl resonances using a 3D NCO-1/2-ROCSA(0.0329,0.467) experiment. The carbonyl is only one example and in the broader picture the family of ROCSA based sequences described here will allow convenient undistorted measurements of many other highly anisotropic systems. The general approach of interspersed symmetry element blocks over *k* indirect points and *s* scans is broadly applicable to tune the interaction Hamiltonian.

## VI. METHODS

### A. Materials

U-^13^C,^15^N-l-histidine (Cambridge Isotope Laboratories (CIL), Andover, MA) re-crystallized in the anionic *τ* tautomer form^57^ and 1-^13^*C*-glycine (CIL, Andover, MA) were ground with a mortar and pestle and packed into 1.6mm MAS NMR rotors.

U-^13^ C,^15^N-Ubiquitin was prepared as described previously.^62^ About 30mg of the protein was crystallized by a batch method in 70% 2-methyl-2,4-pentanediol (MPD), 22% glycerol, 20 mM sodium citrate buffer, pH 4.1. Needle-like crystals were formed under this condition, as previously reported,^63–65^ and were packed into a 1.9mm MAS rotor by ultracentrifugation^66^ using a commercial packing tool (Giotto Biotech, Italy).

### B. Nuclear Magnetic Resonance Spectroscopy

NMR experiments were performed using a Bruker Avance NEO spectrometer operating at 14.1 (*ω*_0*H*_/2*π* = 600 MHz) using a 1.6mm HFXYD probe (Phoenix NMR, Loveland, CO) or 17.6 T (*ω*_0*H*_/2*π* = 750 MHz) using a 1.9mm HCN probe (Bruker BioSpin, Billerica, MA).

NMR experiments at 600 MHz (U-^13^C,^15^N-l-histidine, and 1-^13^*C*-glycine) were performed with spinning frequencies, *ω_r_*/2*π*, of 20000, 16000, or 10000 ± 10 Hz and standard *π*/2 pulse lengths of 1.65, 2.92, and 3.1 *μ*s for the ^1^H, ^13^C, and ^15^N channels respectively corresponding to *ν*_1_ = 151 kHz (^1^H), 86 kHz (^13^C), 81 kHz (^15^N). The RF pulses for the *f*ROCSA(0.0329, 0.467) periods were calibrated to *ν*_1_ = 86, 67, or 43 kHz. The RF pulses for the B block of the *f*ROCSA(0,0.5) experiment were calibrated to *ν*_1_ = 80, 64, or 40 kHz. Two-dimensional *f*ROCSA(*a*_2_, *b*_2_) experiments were taken with between 16 and 256 scans and between 50 and 200 t_1_ increments with an increment time of *nτ_r_*. Continuous-wave ^1^H (*ν*_1_ = 151 kHz) and ^15^N (*ν*_1_ = 25 kHz) decoupling were used during the *f*ROCSA period and ^1^H *ν*_1_ = 151 kHz SWf-TPPM^67^ decoupling during acquisition.

NMR experiments at 750 MHz (U-^13^C,^15^N-Ubiquitin) were performed with a spinning frequency, *ω_r_*/2*π*, of 16666 ± 10 Hz and standard *π*/2 pulse lengths of 2.0, 3.5, and 4.2 *μs* for the ^1^H, ^13^C, and ^15^N channels respectively. The RF pulses for the *f*ROCSA periods were calibrated to *ν*_1_ = 71 kHz. Specific cross-polarization was used for heteronuclear nitrogen-to-carbon polarization transfer, using the ^15^N *ν*_1_ = 1.5 *ν_r_* and ^13^C *ν*_1_ = 2.5 *ν_r_*^68^. The three-dimensional 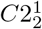 NCO-1/2-ROCSA(0.0329,0.467) experiment was used with 16 scans, 32 t_1_ (*f*ROCSA), and 96 t_2_ (NCO) increments (28 hour total acquisition time). Continuous-wave ^1^H (ν_1_ = 125 kHz) and ^15^N (*ν*_1_ = 25 kHz) decoupling were used during the *f*ROCSA period and ^1^H *ν*_1_ =125 kHz SWf-TPPM^67^ decoupling during acquisition.

Data were processed in Topspin 4.0.7. Additional experimental details and the Bruker pulse programs implemented are given in Figures 1 and S1 and the Supporting Information.

### C. Fitting

To define the experimental anisotropic peaks in the Ubiquitin 3D-*f*ROCSA data set, the peak amplitudes in each isotropic plane were fit with a model containing 58 2D Gaussian peaks. The positions of the peaks (and assignment) were constrained by other 3D NCACX, NCOCX data sets (data not shown). The peak positions were further optimized by fitting the isotropic projections of the 3D (Figure S18).

The experimental anisotropic lineshapes were fit in the frequency domain. The lineshape fitting was done by the POWDER method,^69^ where the powder averaging is performed by computing complete elliptical integrals of the first kind. The resulting lineshapes were convoluted with a Gaussian of unit height, parameterized by its full width at half maximum. As compared to a full numerical simulation, this function could be evaluated quickly, which has allowed us to evaluate parameter uncertainty. Specifically, parameter uncertainties and their correlations were explored via a Markov chain Monte Carlo sampling method. The tabulated errors were determined using the F-test method at 1*σ*.^70^ Experimental sources of uncertainty, such as pulse timings or miscalibrated *ν*_1_ field strengths, may introduce additional uncertainty to the values; however, these uncertainties are not included in the analysis presented here. The errors for Ubiquitin CSA parameters, as reported in the SI, are likely limited by other factors, including isotropic peak overlap, and digital resolution. We think these errors are conservative. The Monte Carlo fitting procedure considerably overestimated the error when compared to errors determined from a more typical non-least square fit routine. The digital resolution was smaller in Ubiquitin than in the small model compound due to differences in processing. We used Bayesian information criterion as a fitting metric to eliminate some of the anisotropic peaks from the analysis (as shown in the SI). These anisotropic peaks likely did not fit well because of lower signal-to-noise or peak overlap. This code used the python packages emcee^71^ and lmfit^70^ for fitting. Nmrglue^72^ was used for accessing the experimental data. The CSA lineshape fitting routines will be made available on GitHub (https://github.com/kfritzsc).

## Supporting information

Supplemental Table: Ubq CSA Values

cp_fROCSA_final.txt

cp_fROCSA_Bprime_final.txt

dbcp_fROCSA_final.txt

## ACKNOWLEDGMENTS

KJF was supported by a Postdoctoral fellowship from the US National Institutes of Health (NIH) (GM128290). EGK was supported by another Postdoctoral fellowship from the NIH (GM135350). AEM was supported by GM088724 from the NIH and 1412253 from the National Science Foundations (NSF). The Ubiquitin NMR data were collected at the New York Structural Biology Center (NYSBC) with support from the Center on Macromolecular Dynamics by NMR Spectroscopy a Biomedical Technology Research Resource NIH (GM118302). The NYSBC is also supported by the Empire State Division of Science Technology and Innovation and Office of Research Infrastructure Program (NIH CO6RR015495). AEM is a member of the NYSBC.

## Notes

### Competing Interest Statement

The authors have declared no competing interest.

## References

1 A. Pines, M. G. Gibby, and J. S. Waugh, Chemical Physics Letters 15, 373 (1972).

2 J. W. Gleeson and R. W. Vaughan, The Journal of Chemical Physics 78, 5384 (1983).

3 H. Sun, L. K. Sanders, and E. Oldfield, Journal of the American Chemical Society 124, 5486 (2002).

4 Y. Wei, A. C. De Dios, and A. E. McDermott, Journal of the American Chemical Society 121, 10389 (1999).

5 Z. Gu, R. Zambrano, and A. McDermott, Journal of the American Chemical Society 116, 6368 (1994).

6 H. Saito, I. Ando, A. Ramamoorthy, S. Hazime, A. Isao, and A. Ramamoorthy, Progress in Nuclear Magnetic Resonance Spectroscopy 57, 181 (2010).

7 B. J. Wylie, C. D. Schwieters, E. Oldfield, and C. M. Rienstra, Journal of the American Chemical Society 131, 985 (2009).

8 J. Kraus, R. Gupta, M. Lu, A. Gronenborn, M. Akke, and T. Polenova, ChemPhysChem, cphc.202000249 (2020).

9 Z. Gu and A. McDermott, Journal of the American Chemical Society 115, 4282 (1993).

10 T. Kameda, N. Takeda, S. Kuroki, H. Kurosu, S. Ando, I. Ando, A. Shoji, and T. Ozaki, Journal of Molecular Structure 384, 17 (1996).

11 R. S. Lipsitz and N. Tjandra, Journal of the American Chemical Society 123, 11065 (2001).

12 P. Allard and T. Hard, Journal of Magnetic Resonance 126, 48 (1997).

13 K. Loth, P. Pelupessy, and G. Bodenhausen, Journal of the American Chemical Society 127, 6062 (2005).

14 P. Schanda and M. Ernst, Progress in Nuclear Magnetic Resonance Spectroscopy 96, 1 (2016).

15 E. G. Keeler, K. J. Fritzsching, and A. E. McDermott, Journal of Magnetic Resonance 296, 130 (2018).

16 E. R. Andrew, A. Bradbury, and R. G. Eades, Nature 182, 1659 (1958).

17 M. A. Alla, E. I. Kundla, and E. T. Lippmaa, JETP Letters 27, 194 (1978).

18 Y. Yarim-Agaev, P. N. Tutunjian, and J. S. Waugh, Journal of Magnetic Resonance (1969) 47, 51 (1982).

19 A. Bax, N. M. Szeverenyi, and G. E. Maciel, Journal of Magnetic Resonance (1969) 51, 400 (1983).

20 R. Tycko, G. Dabbagh, and P. A. Mirau, Journal of Magnetic Resonance (1969) 85, 265 (1989).

21 D. M. Gregory, M. A. Mehta, J. C. Shiels, and G. P. Drobny, Journal of Chemical Physics 107, 28 (1997).

22 S. F. Liu, J. D. Mao, and K. Schmidt-Rohr, Journal of Magnetic Resonance 155, 15 (2002).

23 R. Witter, S. Hesse, and U. Sternberg, Journal of Magnetic Resonance 161, 35 (2003).

24 J. C. Chan and R. Tycko, Journal of Chemical Physics 118, 8378 (2003).

25 R. M. Orr and M. J. Duer, Journal of Magnetic Resonance 181, 1 (2006).

26 Y. Nishiyama, T. Yamazaki, and T. Terao, Journal of Chemical Physics 124, 064304 (2006).

27 J. Herzfeld and A. E. Berger, The Journal of Chemical Physics 73, 6021 (1980).

28 W. T. Dixon, The Journal of Chemical Physics 77, 1800 (1982).

29 A. C. Kolbert and R. G. Griffin, Chemical Physics Letters 166, 87 (1990).

30 Z. Gan, Journal of the American Chemical Society 114, 8307 (1992).

31 J. Z. Hu, D. W. Alderman, C. H. Ye, R. J. Pugmire, and D. M. Grant, Journal of Magnetic Resonance, Series A 105, 82 (1993).

32 O. N. Antzutkin, S. C. Shekar, and M. H. Levitt, TwoDimensional Sideband Separation in Magic-Angle-Spinning NMR (1995).

33 B. J. Wylie, L. J. Sperling, H. L. Frericks, G. J. Shah, W. T. Franks, and C. M. Rienstra, Journal of the American Chemical Society 129, 5318 (2007).

34 A. Bax, N. M. Szeverenyi, and G. E. Maciel, Journal of Magnetic Resonance (1969) 55, 494 (1983).

35 A. Bax, N. M. Szeverenyi, and G. E. Maciel, Journal of Magnetic Resonance (1969) 52, 147 (1983).

36 T. Fujii, T. Onodera, and A. Saika, Chemical Physics Letters 107, 145 (1984).

37 L. Frydman, G. C. Chingas, Y. K. Lee, P. J. Grandinetti, M. A. Eastman, G. A. Barrall, and A. Pines, The Journal of Chemical Physics 97, 4800 (1992).

38 P. J. Grandinetti, Y. K. Lee, J. H. Baltisberger, B. Q. Sun, and A. Pines, Journal of Magnetic Resonance, Series A 102, 195 (1993).

39 M. Carravetta, M. Edén, X. Zhao, A. Brinkmann, and M. H. Levitt, Chemical Physics Letters 321, 205 (2000).

40 B. J. Wylie, W. T. Franks, D. T. Graesser, and C. M. Rienstra, Journal of the American Chemical Society 127, 11946 (2005).

41 B. J. Wylie and C. M. Rienstra, Journal of Chemical Physics 128, 052207 (2008).

42 B. J. Wylie, L. J. Sperling, A. J. Nieuwkoop, W. T. Franks, E. Oldfield, and C. M. Rienstra, Proceedings of the National Academy of Sciences of the United States of America 108, 16974 (2011).

43 D. H. Brouwer and J. A. Ripmeester, Journal of Magnetic Resonance 185, 173 (2007).

44 G. Hou, S. Paramasivam, I.-J. L. Byeon, A. M. Gronenborn, and T. Polenova, Physical Chemistry Chemical Physics 12, 14873 (2010).

45 H. K. Miah, D. A. Bennett, D. Iuga, and J. J. Titman, Journal of Magnetic Resonance 235, 1 (2013).

46 G. Hou, I. J. L. Byeon, J. Ahn, A. M. Gronenborn, and T. Polenova, Journal of Chemical Physics 137, 134201 (2012).

47 M. K. Pandey, M. Malon, A. Ramamoorthy, and Y. Nishiyama, Journal of Magnetic Resonance 250, 45 (2015).

48 D. Mukhopadhyay, C. Gupta, T. Theint, and C. P. Jaroniec, Journal of Magnetic Resonance 297, 152 (2018).

49 M. Fritz, C. M. Quinn, M. Wang, G. Hou, X. Lu, L. M. Koharudin, J. Struppe, D. A. Case, T. Polenova, and A. M. Gronenborn, Physical Chemistry Chemical Physics 20, 9543 (2018).

50 M. Edén and M. H. Levitt, Journal of Chemical Physics 111, 1511 (1999).

51 A. Brinkmann and M. H. Levitt, Journal of Chemical Physics 115, 357 (2001).

52 M. H. Levitt, Journal of Chemical Physics 128, 052205 (2008).

53 M. Hohwy, H. J. Jakobsen, M. Edéen, M. H. Levitt, and N. C. Nielsen, Journal of Chemical Physics 108, 2686 (1998).

54 R. Tycko, Journal of Chemical Physics 126, 64506 (2007).

55 M. J. Potrzebowski, P. Tekely, and Y. Dusausoy, Solid State Nuclear Magnetic Resonance 11, 253 (1998).

56 R. E. Taylor, Concepts in Magnetic Resonance Part A: Bridging Education and Research 22, 79 (2004).

57 S. Li and M. Hong, Journal of the American Chemical Society 133, 1534 (2011).

58 G. Cornilescu and A. Bax, Journal of the American Chemical Society 122, 10143 (2000).

59 F. Cisnetti, K. Loth, P. Pelupessy, and G. Bodenhausen, ChemPhysChem 5, 807 (2004).

60 K. Loth, P. Pelupessy, and G. Bodenhausen, Journal of Biomolecular NMR 27, 159 (2003).

61 R. A. Burton and N. Tjandra, Journal of the American Chemical Society 129, 1321 (2007).

62 A. J. Wand, J. L. Urbauer, R. P. McEvoy, and R. J. Bieber, Biochemistry 35, 6116 (1996).

63 T. I. Igumenova, A. E. McDermott, K. W. Zilm, R. W. Martin, E. K. Paulson, and A. J. Wand, Journal of the American Chemical Society 126, 6720 (2004).

64 T. I. Igumenova, A. J. Wand, and A. E. McDermott, Journal of the American Chemical Society 126, 5323 (2004).

65 K. Y. Huang, G. A. Amodeo, L. Tong, and A. McDermott, Protein Science 20, 630 (2011).

66 G. S. Hisao, M. A. Harland, R. A. Brown, D. A. Berthold, T. E. Wilson, and C. M. Rienstra, Journal of Magnetic Resonance 265, 172 (2016).

67 R. S. Thakur, N. D. Kurur, and P. K. Madhu, Chemical Physics Letters 426, 459 (2006).

68 M. Baldus, A. T. Petkova, J. Herzfeld, and R. G. Griffin, Molecular Physics 95, 1197 (1998).

69 D. W. Alderman, M. S. Solum, and D. M. Grant, The Journal of Chemical Physics 84, 3717 (1986).

70 M. Newville, T. Stensitzki, D. B. Allen, and A. Ingargiola 10.5281/ZENODO.11813 (2014).

71 D. Foreman-Mackey, D. W. Hogg, D. Lang, and J. Goodman, Publications of the Astronomical Society of the Pacific 125, 306 (2013).

72 J. J. Helmus and C. P. Jaroniec, Journal of Biomolecular NMR 55, 355 (2013).

