## Supplemental Information for "Scaled Recoupling of Chemical Shift Anisotropies at High Magnetic Fields under MAS with Interspersed *C*-elements"

(Dated: 2 July 2020)

[a] These authors have contributed equally

### I. TERMINOLOGY AND DEFINITIONS

#### A. Chemical Shift Tensor

This work employs the IUPAC definitions and Haeberlen convention when discussing chemical shift interactions.[1] The isotropic chemical shift is defined as the trace of the chemical shift tensor,

$$\delta_{iso} = \frac{1}{3}(\delta_{zz} + \delta_{yy} + \delta_{xx}), \tag{1}$$

where $\delta_{zz}$, $\delta_{yy}$, and $\delta_{xx}$ are the principal components of the second rank chemical shift tensor. When adopting the Haeberlen convention[1] the principal components are defined as $\mid \delta_{zz} - \delta_{iso} \mid > \mid \delta_{xx} - \delta_{iso} \mid > \mid \delta_{yy} - \delta_{iso} \mid$, and the reduced chemical shift anisotropy, $\delta_{\mathrm{CSA}}$, and asymmetry parameter, $\eta_{\mathrm{CSA}}$ are defined as

$$\delta_{\mathrm{CSA}} = \delta_{zz} - \delta_{iso}, \tag{2}$$

$$\eta_{\mathrm{CSA}} = \frac{\delta_{yy} - \delta_{xx}}{\delta_{\mathrm{CSA}}}. \tag{3}$$

### II. PULSE PROGRAMS

See attached SI_PulsePrograms.zip folder for Bruker pulse programs for the CP-$f$ROCSA (B), CP-$f$ROCSA (B'), and DCP-$f$ROCSA experiments.

### III. SUPPLEMENTARY TABLES AND FIGURES

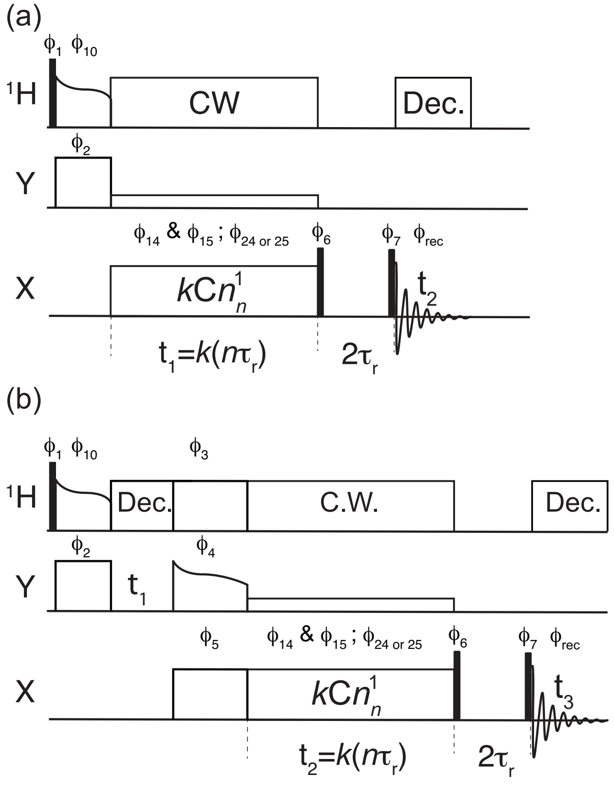


FIG. S 1. Pulse sequence diagrams for the (a) Two-dimensional CP-$f$ROCSA$(a_2, b_2)$ and (b) Three-dimensional DCP-$f$ROCSA$(a_2, b_2)$ experiments.The phase cycle of the CP-$f$ROCSA$(a_2, b_2)$ experiment is: $\phi_1$ = 13, $\phi_{10}$ = 0, $\phi_2$ = 0022, $\phi_{14}$ = 0220, $\phi_{15}$ = 2002, $\phi_{24}$ = 2002, $\phi_{25}$ = 0220, $\phi_6$ = 3113, $\phi_7$ = 0123 2301, $\phi_{rec}$ = 3012 1230 and the DCP-$f$ROCSA$(a_2, b_2)$ experiment is: $\phi_1$ = 13, $\phi_{10}$ = 0, $\phi_2$ = 0, $\phi_3$ = 0, $\phi_4$ = 0, $\phi_5$ = 0022, $\phi_{14}$ = 02, $\phi_{15}$ = 20, $\phi_{24}$ = 2002, $\phi_{25}$ = 0220, $\phi_6$ = 3113, $\phi_7$ = 0123 2301 2301 0123, $\phi_{rec}$ = 3012 1230 1230 3012. Both $\phi_{14}$ and $\phi_{15}$ are used during the A block and either $\phi_{24}$ or $\phi_{25}$ are used during the B (or B') block.

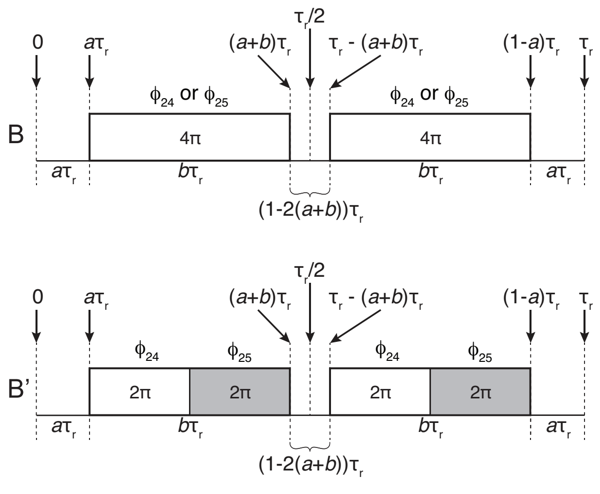


FIG. S 2. Pulse sequence diagram for the B and B' blocks of the *f*ROCSA experiments. The phase cycling of the B' block is given as: $\phi_{24}$ = 2002, $\phi_{25}$ = 0220.

TABLE S I. Fitting results of the carboxyl carbon of the $\alpha$ polymorph of 1-$^{13}$C-glycine

| Experiment | $\delta_{\text{CSA}}$ | $\eta_{\text{CSA}}$ | Theoretical Scaling Factor | Experimental Scaling Factor | Reference |
|---|---|---|---|---|---|
| Non-spinning | -71.1 ± 0.7 ppm | 0.91 ± 0.01 | - | - | This Work |
| ROCSA | -71 ± 3 ppm | 0.89 ± 0.06 | 0.272 | 0.27 ± 0.01 | This Work |
| 7/8-ROCSA(0.0329,0.467) | -72 ± 4 ppm | 0.89 ± 0.07 | 0.249 | 0.25 ± 0.01 | This Work |
| 3/4-ROCSA(0.0329,0.467) | -71 ± 4 ppm | 0.89 ± 0.07 | 0.226 | 0.22 ± 0.01 | This Work |
| 1/2-ROCSA(0.0329,0.467) | -72 ± 5 ppm | 0.88 ± 0.08 | 0.177 | 0.18 ± 0.01 | This Work |
| 1/4-ROCSA(0.0329,0.467) | -72 ± 6 ppm | 0.88 ± 0.09 | 0.130 | 0.13 ± 0.01 | This Work |
| 1/8-ROCSA(0.0329,0.467) | -72 ± 8 ppm | 0.87 ± 0.11 | 0.106 | 0.11 ± 0.01 | This Work |
| MAS | -71.3 ppm | 0.90 | - | - | [2] |
| Non-spinning | -68.3 ppm | 0.95 | - | - | [3] |

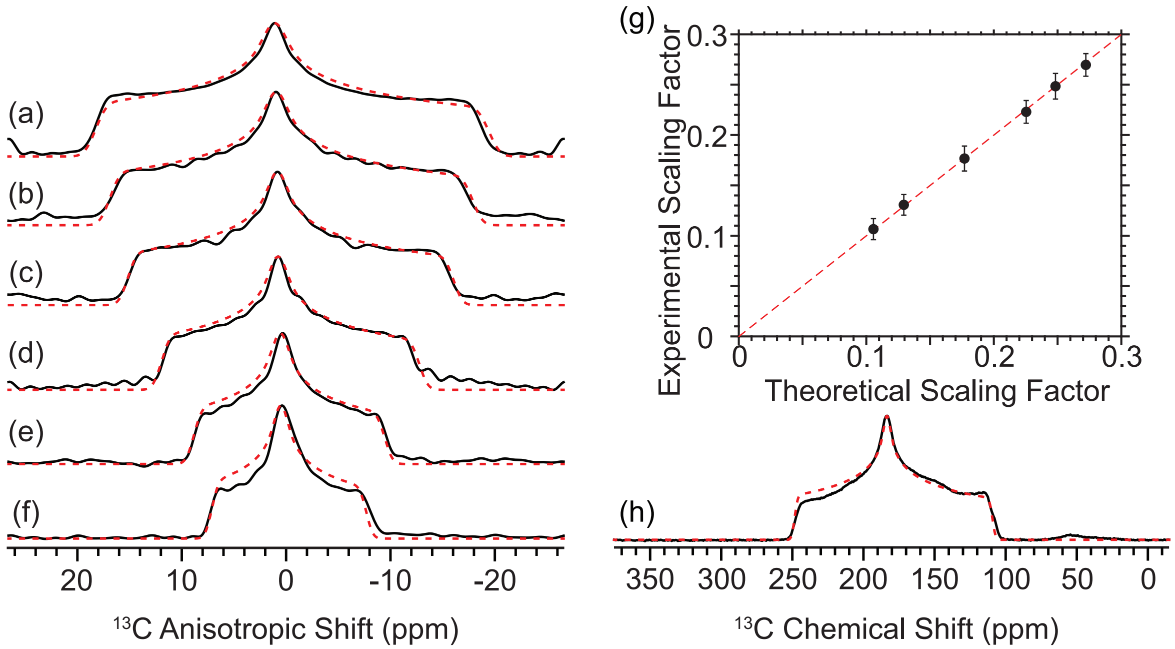


FIG. S 3. (a) $C2_2^1$ ROCSA and (b-f) $C2_2^1$ $f$ROCSA(0.329,0.467) experimental (solid, black) and simulated (dashed, red) lineshapes of the carboxyl carbon of 1-$^{13}$C-glycine for the various sequences described in Figure 1. (g) Plot of the theoretical and experimental scaling factors for the various $f$ROCSA(0.329,0.467) sequences. (h) Experimental (black) and simulated (dashed, red) non-spinning $^{13}$C spectrum of 1-$^{13}$C-glycine. Simulated CSA tensor parameters for each experiment and literature values are given in the Table SI.

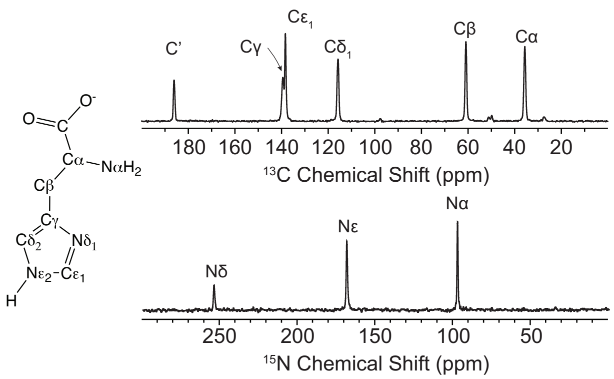


FIG. S 4. $^{13}$C and $^{15}$N CP-MAS spectra and line structure of the $\tau$-tautomer of L-histidine.[4]

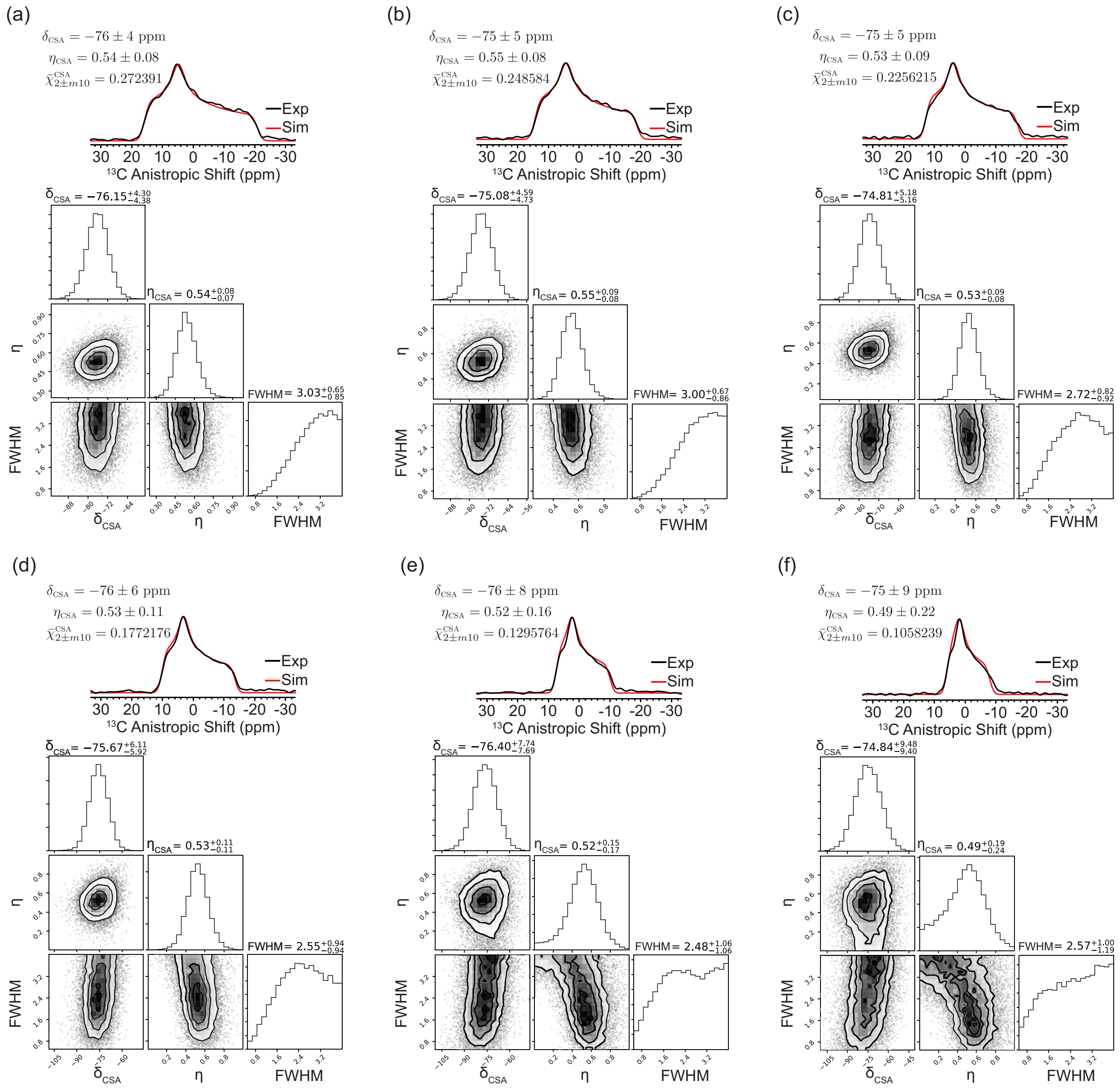


FIG. S 5. (a-f) Fits of the $C2^1_2$ $f$ROCSA(0.0329,0.467) experiments for the carboxyl carbon of L-histidine. The fitting results are shown along with the correlation plots for the various independent variables in the fit and are produced by the python packages emcee[5] and lmfit.[6] The isotropic chemical shift used for the fits was set to carrier frequency of the indirect dimension.

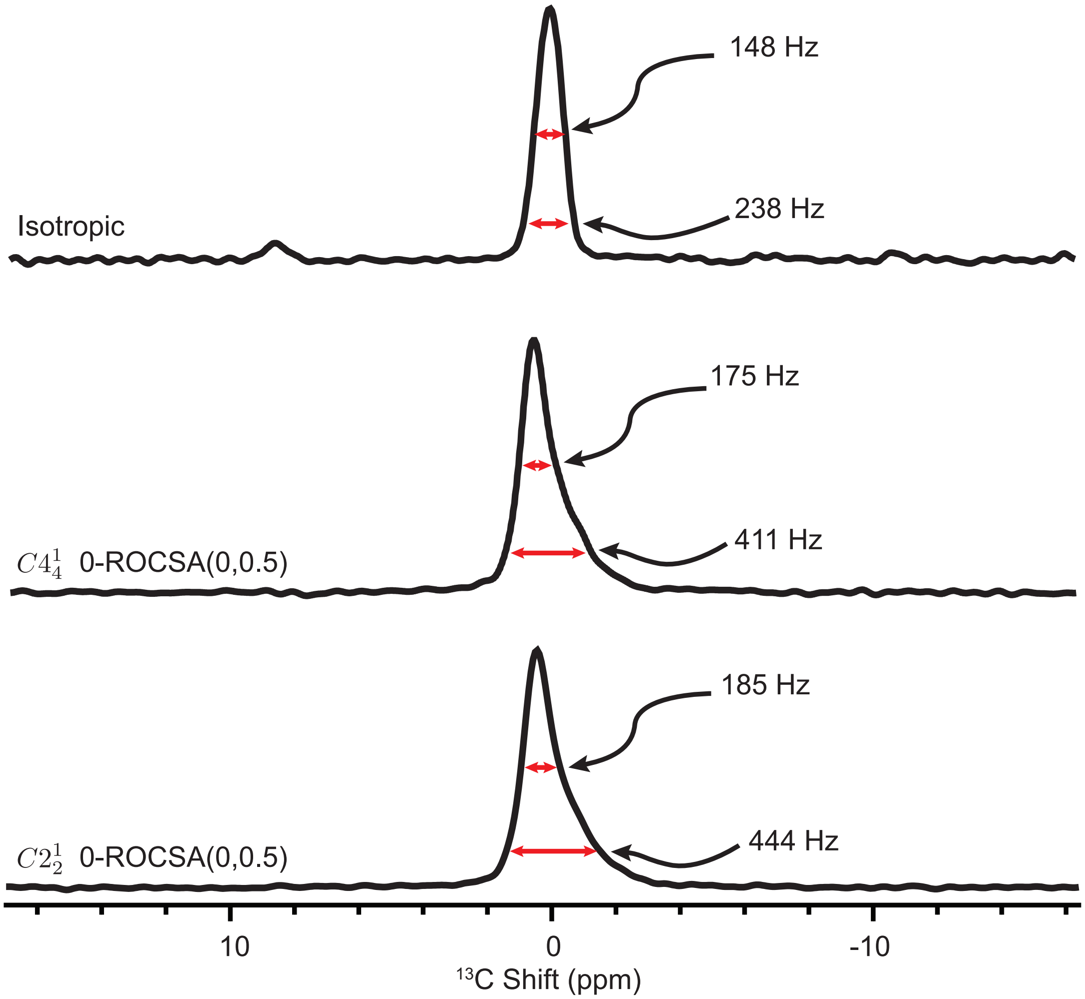


FIG. S 6. Experimental spectra of 0-ROCSA(0,0.5) ($f = 0$ refers to the experiment with only B blocks) experiments using either $C2_2^1$ or $C4_4^1$ symmetry elements. The full width at half maximum (FWHM) and the full width at 15% maximum are given for each spectrum demonstrating a narrowing of the linewidth in the anisotropic dimension when using the $C4_4^1$. The isotropic dimension spectrum is given above showing the linewidth of the isotropic dimension.

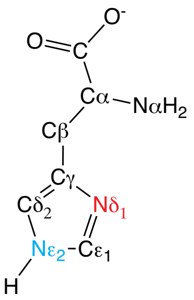


| Site | Experiment | $\delta_{CSA}$ | $\eta_{CSA}$ | Reference |
|---|---|---|---|---|
| $N^{\delta}$ | ROCSA | -229 ± 11 ppm | 0.47 ± 0.06 | This Work |
| $N^{\delta}$ | 3/4-ROCSA | -226 ± 12 ppm | 0.43 ± 0.07 | This Work |
| $N^{\delta}$ | 1/2-ROCSA | -222 ± 14 ppm | 0.39 ± 0.09 | This Work |
| $N^{\epsilon}$ | ROCSA | -114 ± 9 ppm | 0.80 ± 0.11 | This Work |
| $N^{\delta}$ | SUPER | -153 ppm | 0.54 | Ref. 4 |
| $N^{\epsilon}$ | SUPER | -96 ppm | 0.83 | Ref. 4 |

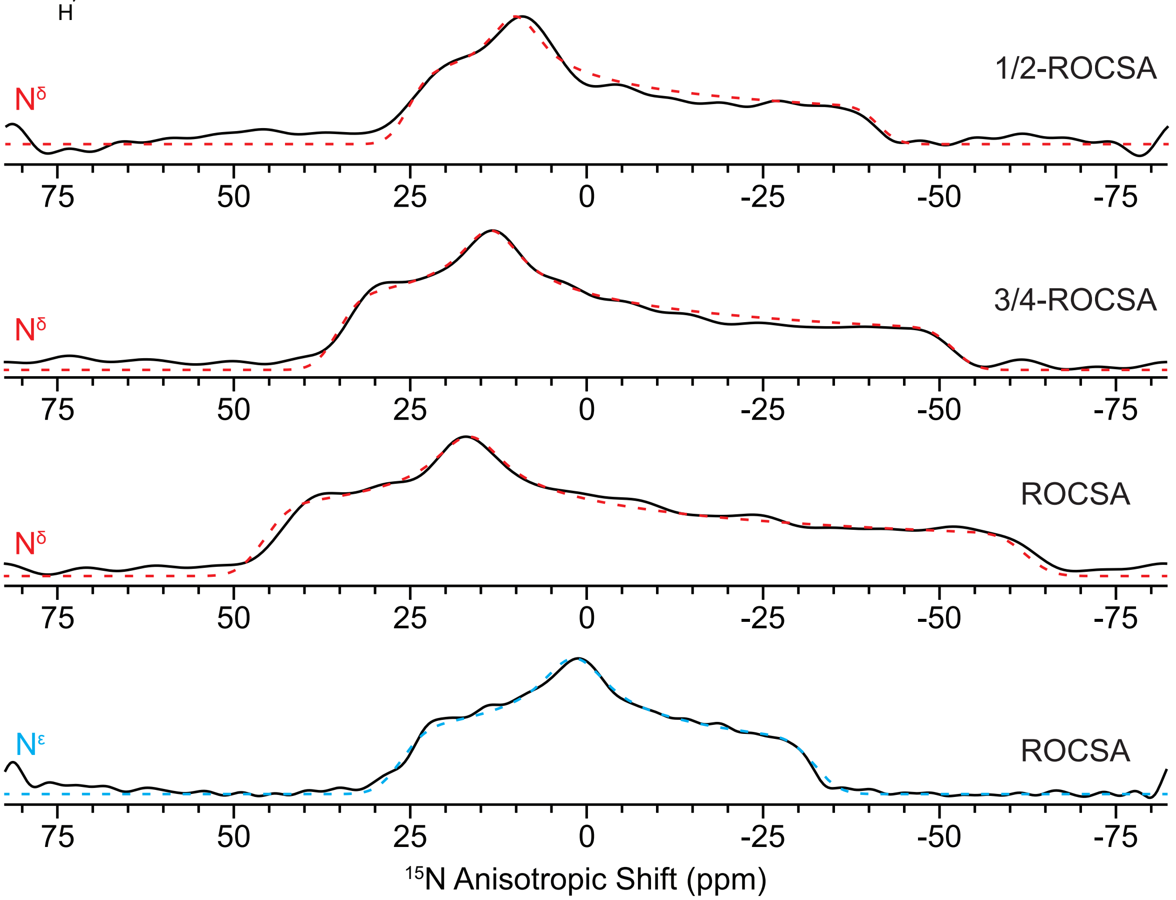


FIG. S 7. Experimental (black) and simulated (dashed) $C2_2^1$ $f$ROCSA(0.0329,0.467) results for the $N^{\delta}$ and $N^{\epsilon}$ nitrogens of L-histidine and the fit CSA tensor parameters, table inset. The line structure of the $\tau$-tautomer of L-histidine is also shown as an inset.[4]

TABLE S II. Fitting results of the carboxyl carbon of the U-$^{13}$C,$^{15}$N-L-histidine at $\nu_r$ = 10 kHz

| Experiment | $\nu_r$ (kHz) | $\delta_{CSA}$ | $\eta_{CSA}$ | $\Omega_{\mathrm{CSA}}$ (ppm) | Spectral Width, Effective (ppm) |
|---|---|---|---|---|---|
| ROCSA | 20 | -76 ± 4 ppm | 0.54 ± 0.08 | ~135 | 244 |
| ROCSA | 12.82 | -74 ± 4 ppm | 0.54 ± 0.06 | ~135 | 156 |
| ROCSA | 12.5 | -81 ± 20 ppm | 0.61 ± 0.10 | ~135 | 153 |
| ROCSA | 10 | -97 ± 18 ppm | 0.62 ± 0.08 | ~135 | 122 |
| 7/8-ROCSA(0.0329,0.467) | 10 | -110 ± 24 ppm | 0.73 ± 0.07 | ~135 | 133 |
| 3/4-ROCSA(0.0329,0.467) | 10 | -75 ± 7 ppm | 0.62 ± 0.08 | ~135 | 147 |
| 1/2-ROCSA(0.0329,0.467) | 10 | -73 ± 4 ppm | 0.53 ± 0.08 | ~135 | 187 |
| 1/4-ROCSA(0.0329,0.467) | 10 | -72 ± 5 ppm | 0.51 ± 0.09 | ~135 | 255 |
| 1/8-ROCSA(0.0329,0.467) | 10 | -71 ± 6 ppm | 0.60 ± 0.10 | ~135 | 313 |

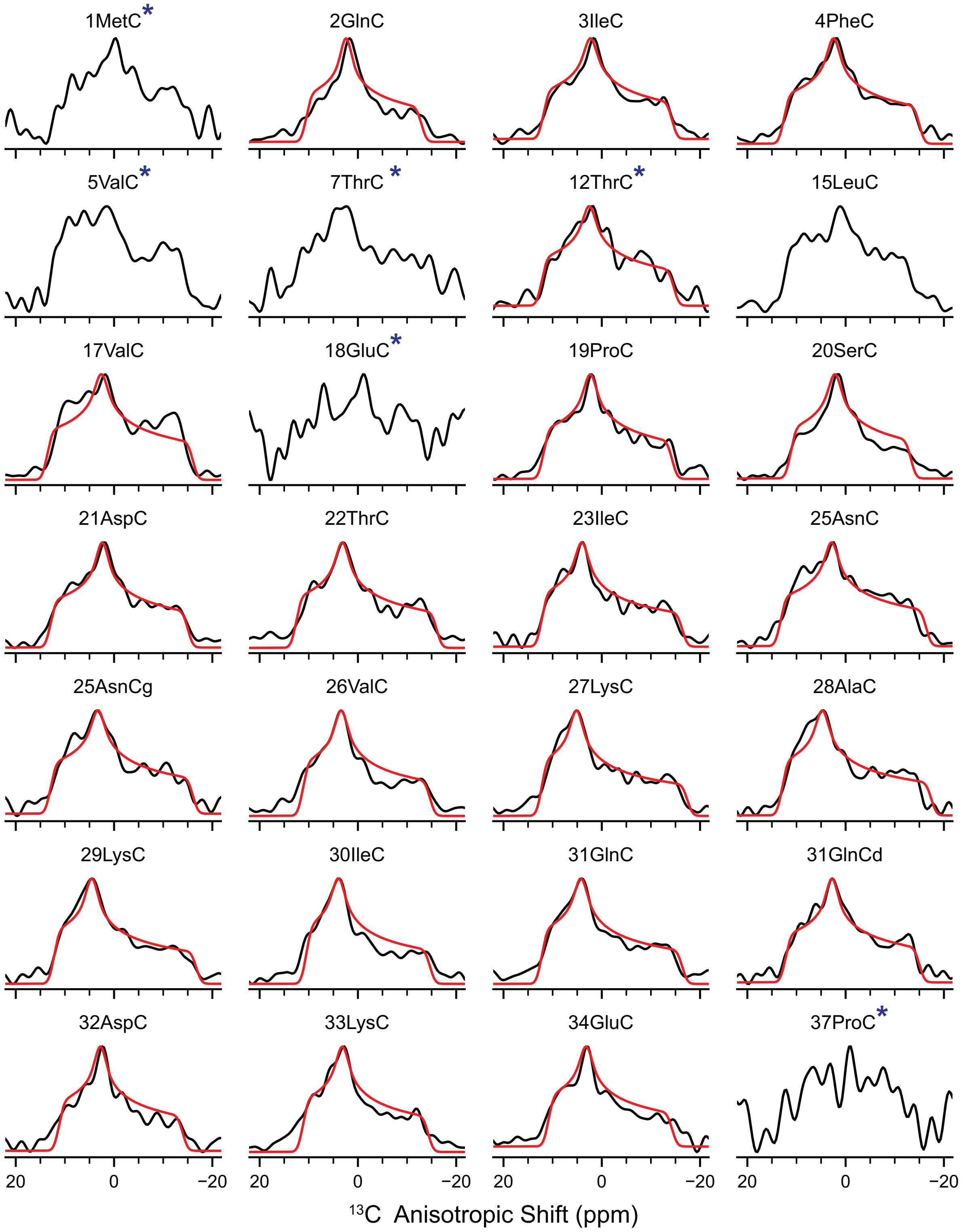


FIG. S 8. Experimental (black) and simulated (red) $C2_2^1$ $f$ROCSA(0.0329,0.467) results for the resolved carbonyls of microcrystalline U-$^{13}$C, $^{15}$N-Ubiquitin. Asterisks indicate peaks that have a signal-to-noise below 25% of the maximal signal-to-noise peak. No fit curve is shown when the BIC fitting metric was larger than -550. See the next two figures for the remaining peaks.

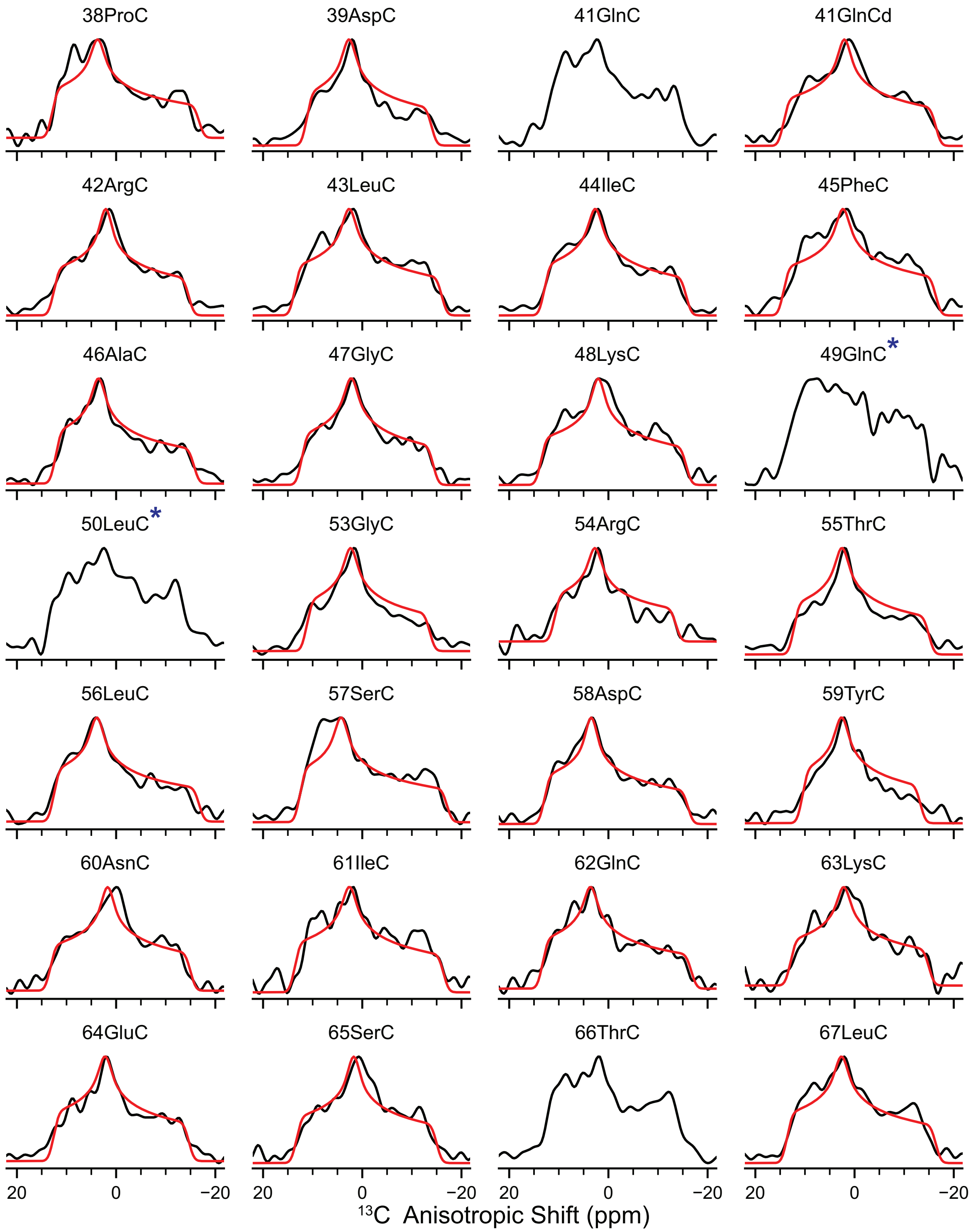


FIG. S 9. Experimental (black) and simulated (red) $C2_2^1$ $f$ROCSA(0.0329,0.467) results for the resolved carbonyls of microcrystalline U-$^{13}$C, $^{15}$N-Ubiquitin. Asterisks indicate peaks that have a signal-to-noise below 25% of the maximal signal-to-noise peak. No fit curve is shown when the BIC fitting metric was larger than -550.

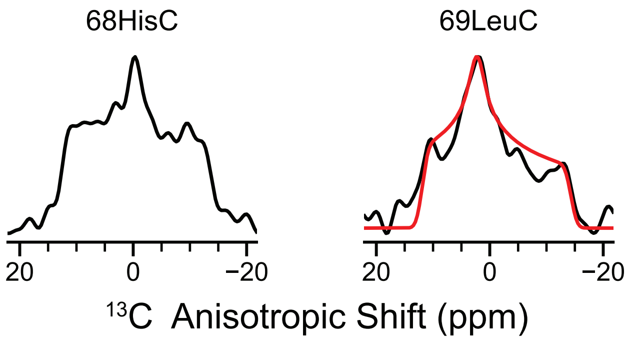


FIG. S 10. Experimental (black) and simulated (red) $C2_2^1$ $f$ROCSA(0.0329,0.467) results for the resolved carbonyls of microcrystalline U-$^{13}$C, $^{15}$N-Ubiquitin. Asterisks indicate peaks that have a signal-to-noise below 25% of the maximal signal-to-noise peak and are excluded from the fitting. No fit curve is shown when the BIC fitting metric was larger than -550.

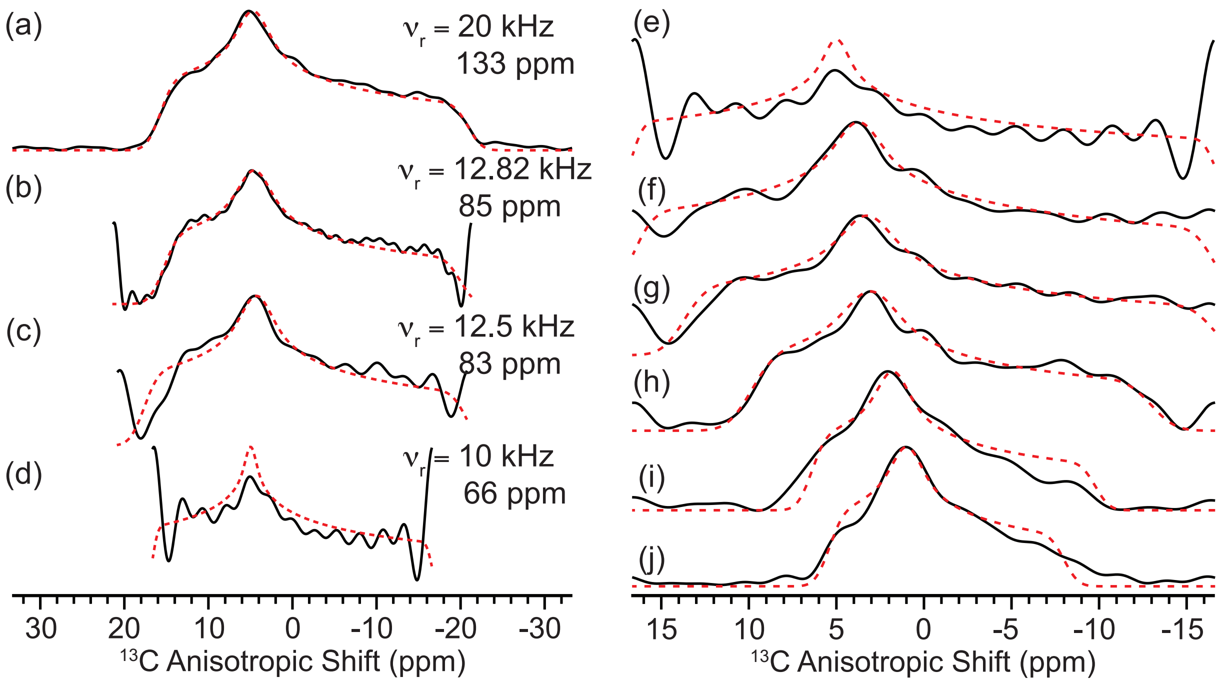


FIG. S 11. (a-e) $C2_2^1$ ROCSA and (f-j) $C2_2^1$ $f$ROCSA(0.329,0.467) experimental (solid, black) and simulated (dashed, red) lineshapes of the carboxyl carbon of U-$^{13}$C,$^{15}$N-L-histidine for the various sequences for different $\nu_r$. (e-j) Demonstrates the family of $f$ROCSA experiments when $\nu_r = 10$ kHz. Simulated CSA tensor parameters for each experiment and effective spectral widths of the experiments are given in Table SII. This demonstrates the difficulty in faithfully extracting the CSA tensor parameters when the effective spectral width of the experiment is greater than the breadth of the lineshape.

| Experiment | $C$-symmetry element | $\delta_{CSA}$ | $\eta_{CSA}$ |
|---|---|---|---|
| ROCSA | $C1_1^1$ | -75 ± 7 ppm | 0.52 ± 0.11 |
| ROCSA | $C2_1^2$ | -75 ± 4 ppm | 0.51 ± 0.08 |
| 1/2-ROCSA(0.0329,0.467) | $C1_1^1$ | -67 ± 9 ppm | 0.58 ± 0.19 |
| 1/2-ROCSA(0.0329,0.467) | $C2_1^2$ | -73 ± 6 ppm | 0.50 ± 0.11 |

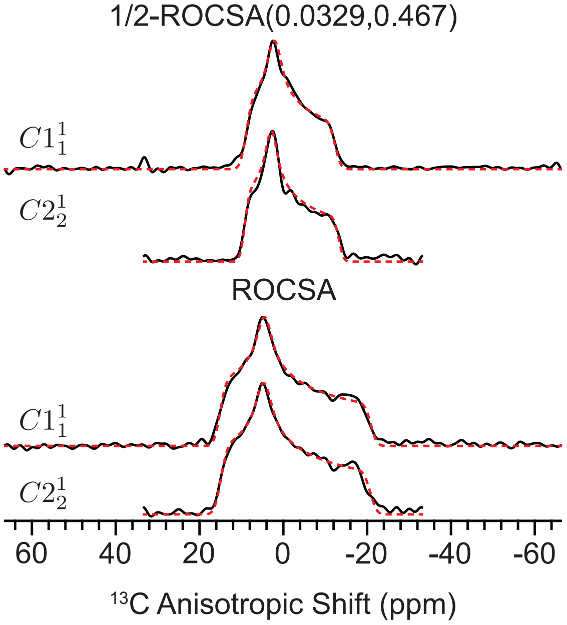


FIG. S 12. Experimental (black) and simulated (red, dashed) $f$ROCSA(0.0329,0.467) results for L-histidine using $C1_1^1$ or $C2_2^1$ symmetry elements. The experiments were acquired with the same indirect acquisition time of 2.5 ms (50 $t_1$ points for $C2_1^2$ and 100 $t_1$ points for $C1_1^1$).

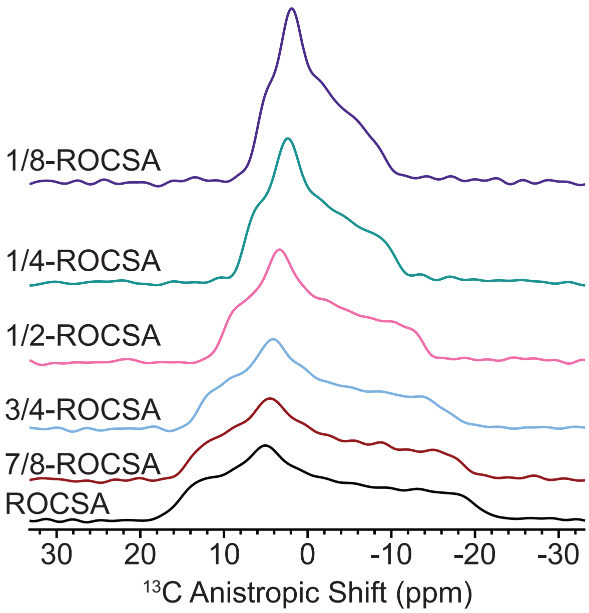


FIG. S 13. Experimental $C2_2^1$ $f$ROCSA(0.0329,0.467) results for the carboxyl carbon of L-histidine scaled to the same absolute scale to demonstrate the increase in signal-to-noise for the versions with smaller scaling factors. These experiments are a fair comparison in spectrometer time, i.e., scans, $t_1$ increments, and processing are identical in each experiment.

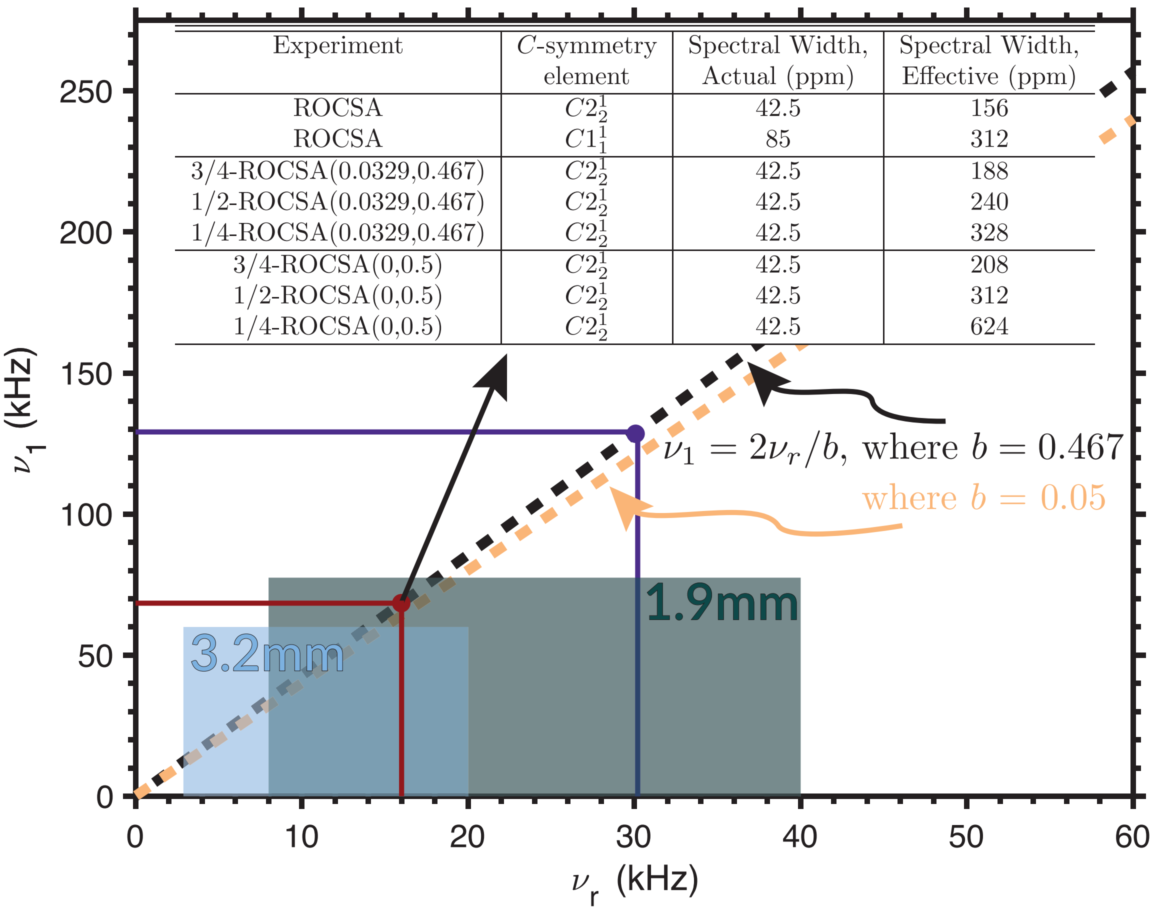


| Experiment | $C$-symmetry element | Spectral Width, Actual (ppm) | Spectral Width, Effective (ppm) |
|---|---|---|---|
| ROCSA | $C2_2^1$ | 42.5 | 156 |
| ROCSA | $C1_1^1$ | 85 | 312 |
| 3/4-ROCSA(0.0329,0.467) | $C2_2^1$ | 42.5 | 188 |
| 1/2-ROCSA(0.0329,0.467) | $C2_2^1$ | 42.5 | 240 |
| 1/4-ROCSA(0.0329,0.467) | $C2_2^1$ | 42.5 | 328 |
| 3/4-ROCSA(0,0.5) | $C2_2^1$ | 42.5 | 208 |
| 1/2-ROCSA(0,0.5) | $C2_2^1$ | 42.5 | 312 |
| 1/4-ROCSA(0,0.5) | $C2_2^1$ | 42.5 | 624 |

FIG. S 14. Plot of the allowable nutation frequency ($\nu_1$) with respect to spinning frequency ($\nu_r$) for the $f$ROCSA$(a_2, b_2)$ experiments at 750 MHz (17.6 T) with the dashed lines, $\nu_1 = 2\nu_r/b$, where $b =$ 0.467 or $= 0.5$. The red (85 ppm) and purple (160 ppm) lines show the desired spinning frequencies for avoiding $R^2$ conditions. Boxes showing the allowable spinning and nutation frequencies ($t > 2$ ms) for standard probes (e.g., 3.2mm HCN EFree, and 1.9mm HCN commercial probes) available for the experiments presented here. Table, inset, of various experiments discussed here and the spectral width considerations of the experiments. The ability to digitize the CSA lineshape for each experiment is an important factor in determining which version of the $f$ROCSA$(a_2, b_2)$ experiments to use ($\Omega_{\text{CSA}} \sim 80\%$ of effective spectral width is recommended). For the 160 ppm spinning frequency condition the spectral widths in the table are multiplied by 1.88.

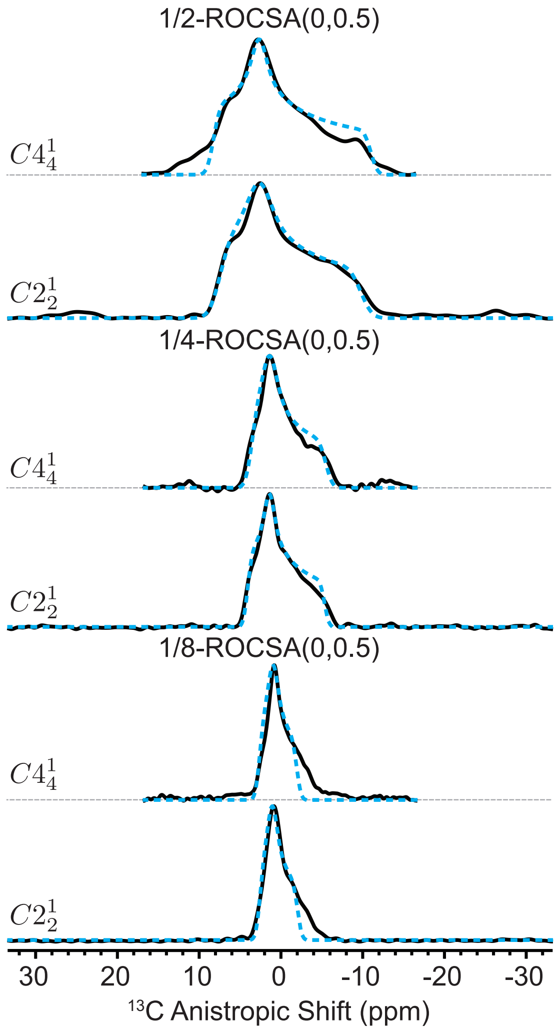


FIG. S 15. Experimental (black) and simulated (cyan, dashed) $f$ROCSA(0,0.5) results for L-histidine using $C2_2^1$ and $C4_4^1$ symmetry elements. The simulated values for the CSA tensors of the $C2_2^1$ and $C4_4^1$ $f$ROCSA(0,0.5) experiments are within error.

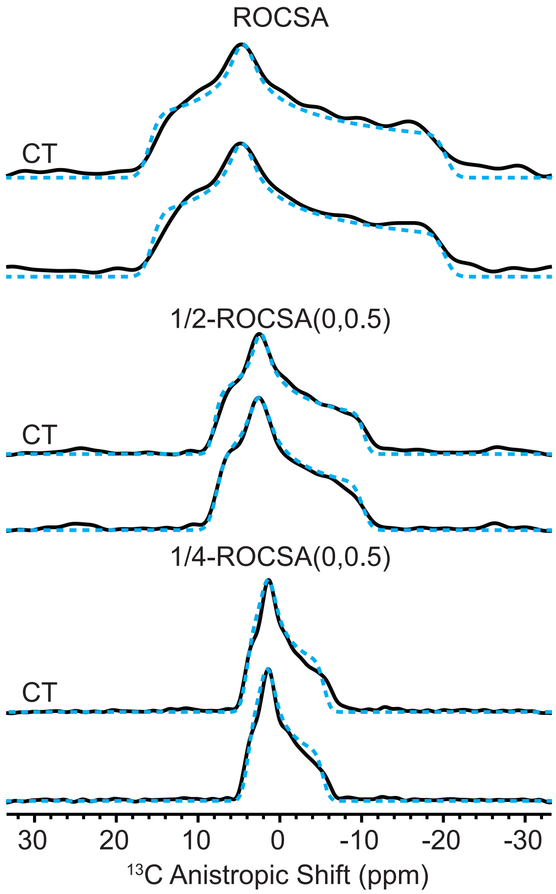


FIG. S 16. Experimental (black) and simulated (cyan, dashed) $C2_2^1$ $f$ROCSA(0,0.5) and $C2_2^1$ CT-$f$ROCSA(0,0.5) results for L-histidine. The simulated values for the CSA tensors for the $f$ROCSA(0,0.5) and CT-$f$ROCSA(0,0.5) experiments are within error.

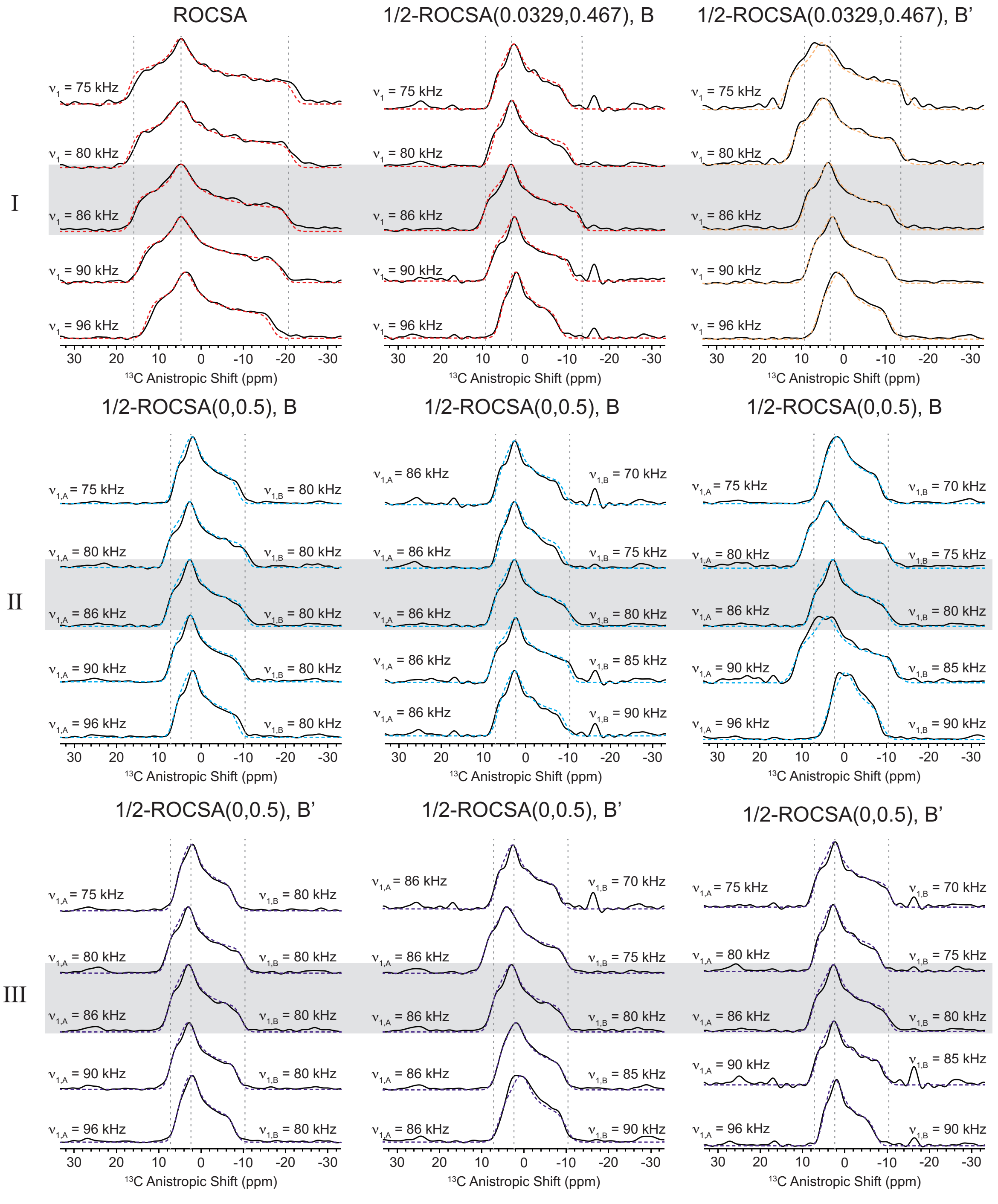


FIG. S 17. Results of the various ROCSA and $f$ROCSA experiments with the various RF field strengths, $\nu_{1,A}$ and $\nu_{1,B}$, miscalibrated by ±5 and ±10 kHz. Results from simulations of the CSA lineshape are given in Tables SIII-V. Vertical dashed gray lines indicate the three principal values $(\delta_{xx}, \delta_{yy}, \delta_{zz})$ of the properly run $f$ROCSA experiment (gray box).

TABLE S III. RF miscalibration results for $f$ROCSA on L-histidine - row I

| Experiment | $\nu_{1,A}$ (kHz) | $\nu_{1,B}$ (kHz) | $\delta_{\text{CSA}}$ | $\eta_{\text{CSA}}$ |
|---|---|---|---|---|
| $C2_2^1$ ROCSA | | | | |
| | 75 | - | -80 ± 5 ppm | 0.55 ± 0.08 |
| | 80 | - | -79 ± 5 ppm | 0.55 ± 0.08 |
| | 86 | - | -76 ± 4 ppm | 0.55 ± 0.08 |
| | 90 | - | -70 ± 4 ppm | 0.53 ± 0.08 |
| | 96 | - | -61 ± 4 ppm | 0.55 ± 0.09 |
| $C2_2^1$ 1/2-ROCSA(0.0329,0.467) - B | | | | |
| | 75 | - | -55 ± 5 ppm | 0.49 ± 0.16 |
| | 80 | - | -61 ± 6 ppm | 0.45 ± 0.14 |
| | 86 | - | -72 ± 6 ppm | 0.50 ± 0.11 |
| | 90 | - | -60 ± 6 ppm | 0.52 ± 0.14 |
| | 96 | - | -45 ± 5 ppm | 0.46 ± 0.21 |
| $C2_2^1$ 1/2-ROCSA(0.0329,0.467) - B' | | | | |
| | 75 | - | -94 ± 9 ppm | 0.52 ± 0.11 |
| | 80 | - | -84 ± 8 ppm | 0.52 ± 0.11 |
| | 86 | - | -74 ± 6 ppm | 0.47 ± 0.11 |
| | 90 | - | -65 ± 8 ppm | 0.50 ± 0.15 |
| | 96 | - | -57 ± 7 ppm | 0.53 ± 0.17 |

TABLE S IV. RF miscalibration results for *f*ROCSA on L-histidine - row II

| Experiment | $\nu_{1,A}$ (kHz) | $\nu_{1,B}$ (kHz) | $\delta_{\text{CSA}}$ | $\eta_{\text{CSA}}$ |
|---|---|---|---|---|
| $C2_2^1$ 1/2-ROCSA(0,0.5) - B | | | | |
| | 75 | 80 | -63 ± 7 ppm | 0.50 ± 0.19 |
| | 80 | 80 | -79 ± 7 ppm | 0.50 ± 0.14 |
| | 86 | 80 | -76 ± 7 ppm | 0.49 ± 0.15 |
| | 90 | 80 | -71 ± 7 ppm | 0.47 ± 0.16 |
| | 96 | 80 | -63 ± 7 ppm | 0.49 ± 0.19 |
| $C2_2^1$ 1/2-ROCSA(0,0.5) - B | | | | |
| | 86 | 70 | -68 ± 7 ppm | 0.45 ± 0.18 |
| | 86 | 75 | -65 ± 7 ppm | 0.42 ± 0.19 |
| | 86 | 80 | -76 ± 7 ppm | 0.49 ± 0.15 |
| | 86 | 85 | -77 ± 7 ppm | 0.53 ± 0.14 |
| | 86 | 90 | -69 ± 7 ppm | 0.45 ± 0.18 |
| $C2_2^1$ 1/2-ROCSA(0,0.5) - B | | | | |
| | 75 | 70 | -63 ± 8 ppm | 0.52 ± 0.20 |
| | 80 | 75 | -88 ± 10 ppm | 0.52 ± 0.14 |
| | 86 | 80 | -76 ± 7 ppm | 0.49 ± 0.15 |
| | 90 | 85 | -100 ± 10 ppm | 0.57 ± 0.13 |
| | 96 | 90 | -53 ± 7 ppm | 0.65 ± 0.22 |

TABLE S V. RF miscalibration results for $f$ROCSA on L-histidine - row III

| Experiment | $\nu_{1,A}$ (kHz) | $\nu_{1,B}$ (kHz) | $\delta_{\text{CSA}}$ | $\eta_{\text{CSA}}$ |
|---|---|---|---|---|
| $C2^1_2$ 1/2-ROCSA(0,0.5) - B' | | | | |
| | 75 | 80 | -60 ± 9 ppm | 0.45 ± 0.22 |
| | 80 | 80 | -77 ± 10 ppm | 0.47 ± 0.17 |
| | 86 | 80 | -74 ± 10 ppm | 0.49 ± 0.18 |
| | 90 | 80 | -69 ± 9 ppm | 0.47 ± 0.19 |
| | 96 | 80 | -60 ± 9 ppm | 0.44 ± 0.23 |
| $C2^1_2$ 1/2-ROCSA(0,0.5) - B' | | | | |
| | 86 | 70 | -66 ± 10 ppm | 0.47 ± 0.21 |
| | 86 | 75 | -82 ± 10 ppm | 0.51 ± 0.15 |
| | 86 | 80 | -74 ± 10 ppm | 0.49 ± 0.18 |
| | 86 | 85 | -68 ± 9 ppm | 0.51 ± 0.19 |
| | 86 | 90 | -66 ± 8 ppm | 0.59 ± 0.19 |
| $C2^1_2$ 1/2-ROCSA(0,0.5) - B' | | | | |
| | 75 | 70 | -77 ± 11 ppm | 0.47 ± 0.18 |
| | 80 | 75 | -65 ± 10 ppm | 0.47 ± 0.21 |
| | 86 | 80 | -74 ± 10 ppm | 0.49 ± 0.18 |
| | 90 | 85 | -70 ± 10 ppm | 0.47 ± 0.20 |
| | 96 | 90 | -54 ± 10 ppm | 0.43 ± 0.24 |

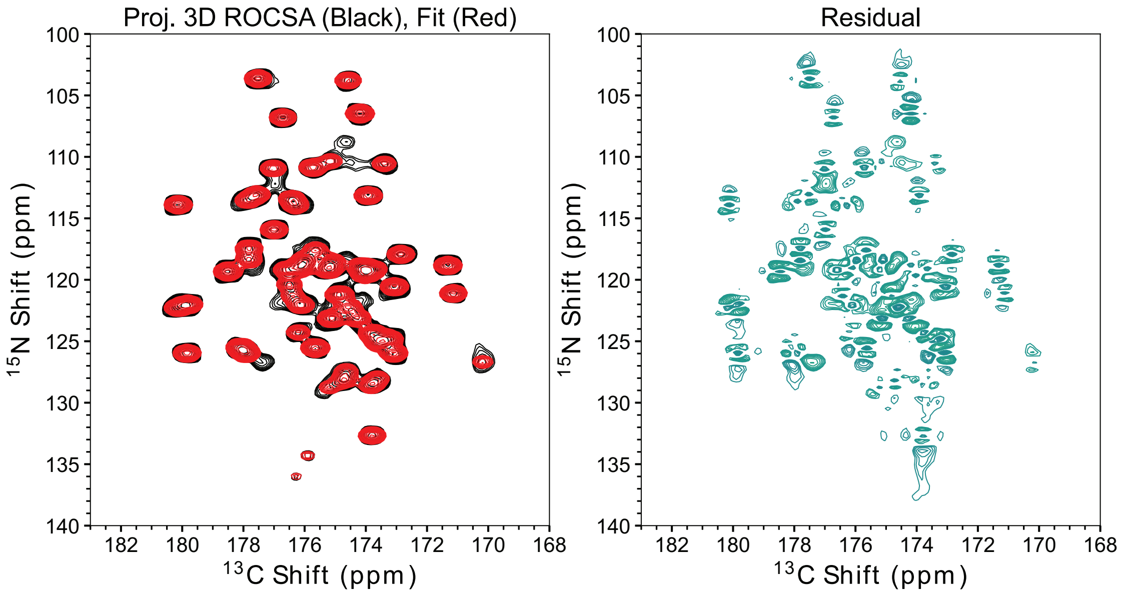


FIG. S 18. Experimental projections of 3D $C2_2^1$ $f$ROCSA(0.0329,0.467) of microcrystalline U-$^{13}$C, $^{15}$N-Ubiquitin (black) and simulated spectrum with 58 2D Gaussian peaks (red). The left shows the fit residual. Note this is the fit of the whole projections, the residual in each slice is approximately 128x less. Most of the residual intensity can be explained by truncation artifacts in the nitrogen dimension. The data and the residual are plotted with a 1.25x multiplier and the lowest contour level of $\pm 4\sigma$ and $\pm 2\sigma$, respectively.

#### REFERENCES


[1] R. K. Harris, E. D. Becker, S. M. Cabral De Menezes, P. Granger, R. E. Hoffman, and K. W. Zilm, Pure and Applied Chemistry **80**, 59 (2008).

[2] M. J. Potrzebowski, P. Tekely, and Y. Dusausoy, Solid State Nuclear Magnetic Resonance **11**, 253 (1998).

[3] R. E. Taylor, Concepts in Magnetic Resonance Part A: Bridging Education and Research **22**, 79 (2004).

[4] S. Li and M. Hong, Journal of the American Chemical Society **133**, 1534 (2011).

[5] D. Foreman-Mackey, D. W. Hogg, D. Lang, and J. Goodman, Publications of the Astronomical Society of the Pacific **125**, 306 (2013).

[6] M. Newville, T. Stensitzki, D. B. Allen, and A. Ingargiola 10.5281/ZENODO.11813 (2014).
