## Supplementary material for "Scaled Recoupling of Chemical Shift Anisotropies at High Magnetic Fields under MAS with Interspersed *C*-elements": cp_fROCSA_final.txt

;F1{F2}CP F1 fROCSA
;All fROCSA possible
;7/8 >= f >= 1/8 possible
;Cn^1_n; n=2 only
;Avance III+NEO, TopSpin 3.2+TopSpin 4.0.7, KJF+EGK 20200702

;*********************************************************
;WARNING: This sequence allows for multiple versions of
;fROCSA to be performed and therefore requires the proper
;setting of multiple loops and constants for the exp. to
;run correctly. READ CAREFULLY!
;*********************************************************

;d1 : recycle delay
;p1 : F1 90
;p2 : F2 90
;p20 : CP contact time
;pl1 : F1 power @ p1
;pl2 : F2 power @ p2
;pl21 : F1 CP power
;pl22 : F2 CP power (nominal)
;p11 : F1 ROCSA 2 Pi Rotation.
;pl11 : F1 ROCSA power for p11
;sp22 : H->F1 CP 1H power shaped
;pl12 : 1H  decoupling (normal)
;pl13 : 1H  decoupling (other)
;pl14 : Y decoupling during ROCSA (e.g., cw_14)
;pcpd2 : 1H pulse length in decoupling sequence
;cpdprg2 : 1H decoupling sequence for acquisition
;cpdprg3 : 1H decoupling sequence during fROCSA
;cpdprg4 : 15N decoupling sequence during fROCSA
;cnst31 : MAS spinning speed in HZ
;cnst32 : a1 = 0.0329 (Chan and Tycko 2003) - A block
;cnst33 : b1 = 0.4670 (Chan and Tycko 2003) - A block
;cnst34 : a2= B block timing
;cnst35 : b2= B block timing
;l29 : = 1 for FullROCSA, 2 for 1/2, 4 for 1/4 or 3/4, 8 for 1/8 or 7/8
;l26 : = 0 for 1/2,1/4,1/8 and >0 for 1/2,3/4,7/8
;l30 : number of rotor periods for z-filter (=2,3,4)
;cnst28 : set to 0: counter for number of times through A/B loops
;cnst29 : set to 0: counter for number of times through A/B loops
;NS : MUST be = 16*n
;FnMODE = STATES-TPPI
;REVERSE = TRUE for indirect dimension for correct direction

;set acquisition for baseopt mode
"acqt0=-p1/2"
;set CP shape pulse to 1/4 of set power for tangential pulse
"spw22=plw22/0.25"

;set d0 to 3u for first loop
"d0=3u"
;set l0 loop to 0 for first loop - no fROCSA in 1st loop
"l0=0"
;set loop counters for phase cycle reset to zero
"cnst28=0"
"cnst29=0"

;define delays for the fROCSA loops
define delay datr
define delay dbtr
define delay dctr
define delay ddtr
define delay tr
"tr = (1s/cnst31)"
"datr= cnst32 * tr"
"dbtr= tr/2 - (cnst32+cnst33)*tr"
"dctr= cnst34 * tr"
"ddtr= tr/2 - (cnst34+cnst35)*tr"

;Calculate the 2pi pulses for fROCSA
"p11= cnst33*(1s/(2*cnst31))"
"p9= cnst35*(1s/(2*cnst31))"

;Calculate the delay for XY-16 decoupling if used (xy_16.egk for fROCSA - 15N)
"d17=tr-p17"

;Calculate the Z-filter delay
define delay dzfilt
"dzfilt = l30*(1s/cnst31)-0.1u-p1"

1 ze

;****************************************************************
;Power level, pulse length and recycling delay protection modules
;****************************************************************

if "(p2+p20+p30+td*dw)>50m" {
1u
print "pulse length exceeds limit!"
goto HaltAcqu
}

;***********************************************************************
;END of power level, pulse length and recycling delay protection modules
;***********************************************************************

	1u pl22:f2

2 d1 do:f2

; CP
  (p2 pl2 ph1):f2
  (p20 pl21 ph2):f1 (p20:sp22 ph10):f2
 	0.1u cpds3:f2 cpds4:f3

if "l26==0"
{ ; fROCSA 1/2,1/4,1/8, or 1
	if "l0==0"
	{
	}
	else
	{
;		ROCSA Cn^1_n
	5
		if "cnst28%l29==0"
			{
; 		A
  6	datr
  		(p11*0.25 pl11 ph14):f1
  		(p11 pl11 ph15):f1
  		(p11*0.75 pl11 ph14):f1
  		dbtr
			;2nd-half
  		dbtr
  		(p11*0.75 pl11 ph14):f1
			(p11 pl11 ph15^):f1
			(p11*0.25 pl11 ph14^):f1
			datr
			lo to 6 times 2
			}
		else
			{
;			B-block
	7		dctr
  		(p9*0.25 pl9 ph24):f1
  		(p9 pl9 ph24):f1
  		(p9*0.75 pl9 ph24):f1
  		ddtr
			;2nd-half
  		ddtr
  		(p9*0.75 pl9 ph24):f1
			(p9 pl9 ph24):f1
			(p9*0.25 pl9 ph24^):f1
			dctr
			lo to 7 times 2
			}
	"cnst28=cnst28+1"
			if "cnst28==l29*2"
				{
					"cnst28=0"
					0.1u rpp14 rpp15 rpp24
				}
			else
				{
				}
	lo to 5 times l0
	}
}
else
{ ; fROCSA 1/2,3/4,7/8,1
	if "l0==0"
		{
		}
	else
		{
;			ROCSA Cn^1_n
		8
		if "cnst29%l29==0"
			{
;			B-block
		9 dctr
  		(p9*0.25 pl9 ph25):f1
  		(p9 pl9 ph25):f1
		  (p9*0.75 pl9 ph25):f1
		  ddtr
			;2nd-half
		  ddtr
 		  (p9*0.75 pl9 ph25):f1
			(p9 pl9 ph25):f1
			(p9*0.25 pl9 ph25^):f1
			dctr
			lo to 9 times 2
			}
		else
			{
;			A-block
	10  datr
		  (p11*0.25 pl11 ph14):f1
		  (p11 pl11 ph15):f1
		  (p11*0.75 pl11 ph14):f1
		  dbtr
			;2nd-half
		  dbtr
		  (p11*0.75 pl11 ph14):f1
			(p11 pl11 ph15^):f1
			(p11*0.25 pl11 ph14^):f1
			datr
			lo to 10 times 2
			}
	"cnst29=cnst29+1"
			if "cnst29==l29*2"
				{
					"cnst29=0"
					0.1u rpp14 rpp15 rpp25
				}
			else
				{
				}
	lo to 8 times l0
		}
}

(p1 pl1 ph5):f1
0.1u do:f2 do:f3
dzfilt
(p1 pl1 ph6):f1

  go=2 ph31 cpds2:f2 finally do:f2
	0.1u rpp14 rpp15 rpp24 rpp25
	"cnst28=0"
	"cnst29=0"
  10m mc #0 to 2 F1PH(calph(ph5,+90), calclc(l0,1))
HaltAcqu, 1m
11 exit

ph0= 0
ph1= 1 3
ph10= 0
ph2= 0 0 2 2; MC2= 0 2 2 0

ph14= 0 2 2 0 0 2 2 0
ph15= 2 0 0 2 2 0 0 2

ph24= 2 0 0 2

ph25= 0 2 2 0

ph5= 3 1 1 3; M = + + + +

ph6 = 0 1 2 3 2 3 0 1; M = 3 0 1 2 1 2 3 0
			2 3 0 1 0 1 2 3

ph31= 3 0 1 2 1 2 3 0
			1 2 3 0 3 0 1 2
