## Supplementary material for "Scaled Recoupling of Chemical Shift Anisotropies at High Magnetic Fields under MAS with Interspersed *C*-elements": dbcp_fROCSA_final.txt

;F1{F2}DoubleCP F1 fROCSA
;All fROCSA possible
;7/8 >= f >= 1/8 possible
;Cn^1_n; n=2 only
;Avance III+NEO, TopSpin 3.2+TopSpin 4.0.7, KJF+EGK 20200702

;acqt0 parameter is optimized to have ph1= 0; fcor = 0.5
;d1 : recycle delay

;********CARBON*******

;p30 : 15N -> 13C Specific-CP pulse
;pl1 : NOT USED
;sp31 : 15N -> 13C Specific-CP power level (use only tan-8-2-60.and)

;********PROTON*********

;p2 : 1H 90 deg pulse at pl2
;p20 : 1H-15N CP time
;spnam20 : 1H CP shape pulse @ p20 (use only tan-8-2-60.and)
;cpdprg2 : sequence used for decoupling (spinal64 recommended)
;pcpd2 : pulse length in decoupling sequence
;cpdprg3 : 1H decoupling sequence during fROCSA

;pl2 : 1H 90 @ p2
;pl12 : 1H decoupling
;sp22 : 1H CP power @ p20 (use tan-8-2-60.and only)
;pl32 : 1H decoupling during Specific-CP

;********NITROGEN*******

;pl3 : not used
;pl23 : 15N CP power @ p20
;pl33 : 15N DCP power @ p30
;cnst3 : 15N carrier offset for DCP transfer, usually around 0ppm
;cpdprg4 : 15N decoupling sequence during fROCSA

;*********fROCSA***********
;pl13 : 1H  decoupling (other)
;pl14 : Y decoupling during ROCSA (xy_16.egk)
;p17 : Y pi pulse for xy decoupling
;d17 : rotor sync delay for xy decoupling (tr-p17)
;pcpd2 : 1H pulse length in decoupling sequence
;cpdprg2 : decoupling sequence
;cnst31 : MAS spinning speed in HZ
;cnst32 : a1 = 0.0329 (Chan and Tycko 2003) - A block
;cnst33 : b1 = 0.4670 (Chan and Tycko 2003) - A block
;cnst34 : a2= B block timing
;cnst35 : b2= B block timing
;l29 : = 1 for FullROCSA, 2 for 1/2, 4 for 1/4 or 3/4, 8 for 1/8 or 7/8
;cnst28 : set to 0: counter for number of times through A/B loops
;l26 : =0 for 1/2,1/4,1/8 and >0 for 1/2,3/4,7/8
;NS : MUST be = 16*n
;FnMODE = STATES-TPPI
;REVERSE = TRUE for indirect dimension for correct direction
;F3 : direct dimension
;F2 : fROCSA dimension
;F1 : indirect Specific-CP dimension


#include <Avancesolids.incl>


;setting up actual CP shape pulse power level to adjust shaped pulse to 100%
;average power level

;"spw22=plw22/(integfac22*integfac22)"
;"spw31=plw31/(integfac31*integfac31)"

"acqt0=(-p2/2)"
"in0=inf2"
"d0=1u"

"in0=inf1"
;set l0 loop to 0 for first loop - no fROCSA in 1st loop
"l0=0"
;set loop counters for phase cycle reset to zero
"cnst28=0"
"cnst29=0"

;define delays for the fROCSA loops
define delay datr
define delay dbtr
define delay dctr
define delay ddtr
define delay tr
"tr = (1s/cnst31)"
"datr= cnst32 * tr"
"dbtr= tr/2 - (cnst32+cnst33)*tr"
"dctr= cnst34 * tr"
"ddtr= tr/2 - (cnst34+cnst35)*tr"

;Calculate the 2pi pulses for fROCSA
"p11= cnst33*(1s/(2*cnst31))"
"p9= cnst35*(1s/(2*cnst31))"

;Calculate the delay for XY-16 decoupling if used (xy_16.egk for fROCSA - 15N)
"d17=tr-p17"

;Calculate the Z-filter delay
define delay dzfilt
"dzfilt = 2*(1s/cnst31)-0.1u-p1"


1 ze

2 d1 do:f2

;****************************************************************
;Power level, pulse length and recycling delay protection modules
;****************************************************************

if "(p2+p20+p30+td*dw)>50m" {
1u
print "pulse length exceeds limit!"
goto HaltAcqu
}

;***********************************************************************
;END of power level, pulse length and recycling delay protection modules
;***********************************************************************

; CP


  (p2 pl2 ph1):f2
  (p20 pl23 ph2):f3 (p20:sp22 ph10):f2

if "l1==1"{
"d0=in0-3u"
}
if "l1>0"{
  0.1u cpds2:f2
  (center (d0) (p1*2 pl1 ph8):f1)   ; refocus 13C J
  0.1u do:f2
}

; Specific-CP
   1u fq=cnst1(sfo ppm):f1; Move F1 on res in ppm from SFO1
   1u fq=cnst3(sfo ppm):f3; Move F1 on res in ppm from SFO1
  (p30:sp31 ph4):f1 (p30 pl32 ph3):f2  (p30 pl33 ph5):f3 ;DCP 15N-13C

	0.1u cpds3:f2 cpds4:f3
  1u fq=0:f1; Move 13C to center

if "l26==0"
{ ; fROCSA 1/2,1/4,1/8, or 1
	if "l0==0"
	{
	}
	else
	{
;		ROCSA Cn^1_n
	5
		if "cnst28%l29==0"
			{
; 		A
  6	datr
  		(p11*0.25 pl11 ph14):f1
  		(p11 pl11 ph15):f1
  		(p11*0.75 pl11 ph14):f1
  		dbtr
			;POST
  		dbtr
  		(p11*0.75 pl11 ph14):f1
			(p11 pl11 ph15):f1
			(p11*0.25 pl11 ph14):f1
			datr ipp14 ipp15
			lo to 6 times 2
			}
		else
			{
;			B
;			0.1u
;			print "B-cycle"
	7		dctr
  		(p9*0.25 pl9 ph24):f1
  		(p9 pl9 ph25):f1
  		(p9*0.75 pl9 ph24):f1
  		ddtr
			;POST
  		ddtr
  		(p9*0.75 pl9 ph24):f1
			(p9 pl9 ph25):f1
			(p9*0.25 pl9 ph24):f1
			dctr ipp24 ipp25
			lo to 7 times 2
			}
	"cnst28=cnst28+1"
	lo to 5 times l0
	}
}
else
{ ; fROCSA 1/2,3/4,7/8,1
	if "l0==0"
		{
		}
	else
		{
;			ROCSA Cn^1_n
		8
		if "cnst29%l29==0"
			{
;			B
;			0.1u
;			print "B-cycle"
		9 dctr
  		(p9*0.25 pl9 ph24):f1
  		(p9 pl9 ph25):f1
		  (p9*0.75 pl9 ph24):f1
		  ddtr
			;POST
		  ddtr
 		  (p9*0.75 pl9 ph24):f1
			(p9 pl9 ph25):f1
			(p9*0.25 pl9 ph24):f1
			dctr ipp24 ipp25
			lo to 9 times 2
			}
		else
			{
;			A
	10  datr
		  (p11*0.25 pl11 ph14):f1
		  (p11 pl11 ph15):f1
		  (p11*0.75 pl11 ph14):f1
		  dbtr
			;POST
		  dbtr
		  (p11*0.75 pl11 ph14):f1
			(p11 pl11 ph15):f1
			(p11*0.25 pl11 ph14):f1
			datr ipp14 ipp15
			lo to 10 times 2
			}
	"cnst29=cnst29+1"
	lo to 8 times l0
		}
}

(p1 pl1 ph6):f1
0.1u do:f2 do:f3
dzfilt
(p1 pl1 ph7):f1

  go=2 ph31 cpds2:f2 finally do:f2
	0.1u rpp14 rpp15 rpp24 rpp25
	"cnst28=0"
	"cnst29=0"
  10m mc #0 to 2
	F1PH(calph(ph6,+90), calclc(l0,1))
	F2PH(calph(ph2, +90), caldel(d0, +in0) & calclc(l1, 1))
  HaltAcqu, 1m
4 exit

ph0 = 0
ph10 = 0
ph1 = 1 3
ph2 = 0
ph3 = 0
ph4 = 0
ph8 = 1
ph5 = 0 0 2 2; MC2= 0 2 2 0

ph14= 0 2 2 0 0 2 2 0
ph15= 2 0 0 2 2 0 0 2

ph24= 2 0 0 2

ph25= 0 2 2 0

ph6= 3 1 1 3; M = + + + +

ph7 = 0 1 2 3 2 3 0 1; M = 3 2 1 0 1 0 3 2
      2 3 0 1 0 1 2 3

ph31= 3 0 1 2 1 2 3 0
			1 2 3 0 3 0 1 2
